# A super-pangenome of *Capsicum* reveals genomic basis of domestication, pungency, and metabolic diversity

**DOI:** 10.64898/2026.11.17.767367

**Authors:** Jiantao Zhao, Cheng Xiong, Xuebo Zhao, Pei Zhao, Honghe Sun, Lujia Yu, Yi Liu, Robert Jarret, Fan Zhu, Xiongze Dai, Sha Yang, Zhoubin Liu, Huiyang Yu, Lijun Ou, Bozhi Yang, Hao Xu, Yanlong Li, Zhongyi Wang, Bowen Hu, Rui Weng, Xuexiao Zou, Zhangjun Fei, Feng Liu

## Abstract

The *Capsicum* genus encompasses remarkable genetic and phenotypic diversity underlying its agricultural, economic, and cultural importance. Here, we present a *Capsicum* super-pangenome constructed from 48 chromosome-scale genomes, including 38 newly assembled, spanning all five domesticated taxa and their wild relatives. We identified 72,687 pangene families and abundant structural variants (SVs), revealing lineage-specific gene-content dynamics and patterns consistent with strong purifying selection against large SVs. A pan-NLRome revealed extensive presence-absence variation in the disease-resistance gene repertoire, with domesticated accessions harboring fewer NLRs than their wild progenitors. The graph-based super-pangenome enabled extensive genome-wide association mapping and robust genomic prediction across 130 agronomic and metabolic traits, leading to the identification and functional validation of *RAP2* as a regulator of carotenoid accumulation and the discovery of a structurally complex terpene synthase locus associated with sesquiterpene ylangene. Pangenome-wide dissection of capsaicinoid biosynthesis further revealed evolutionary remodeling of the pathway through tandem gene duplication, interchromosomal rearrangement, and regulatory specialization. Collectively, our findings illuminate the genomic basis of *Capsicum* domestication and metabolic diversity and establish a comprehensive resource for genomics-assisted improvement of disease resistance, fruit quality, and flavor.

## Introduction

Pepper (*Capsicum* spp.) belongs to the genus *Capsicum*, which comprises approximately 35 species, including five domesticated taxa: *C. annuum* var. *annuum*, *C. chinense*, *C. frutescens*, *C. baccatum* var. *pendulum*, and *C. pubescens*. Originating in the Americas (Kim et al., 2014), peppers are now among the most widely cultivated crops worldwide and have profoundly shaped diets and culinary traditions across diverse cultures (Katz, 2019). They also hold substantial economic and cultural importance, with diverse applications in cuisine, medicine, and industry (Hernández-Pérez et al., 2020). Genomic studies have begun to reveal how genetic diversity and gene flow have shaped pepper domestication and population differentiation (Liu et al., 2023). However, the genetic basis of key agronomic traits, including disease resistance, fruit morphology, nutritional quality, and volatile composition, remains largely unexplored, particularly in underrepresented domesticated taxa such as *C. pubescens* and *C. baccatum*. Addressing these knowledge gaps requires a comprehensive framework capable of capturing the full spectrum of genomic variation across *Capsicum* species and linking this variation to agronomic and metabolomic diversity.

Although more than 20 pepper genome assemblies have been published, many were generated using short-read or earlier long-read sequencing technologies with lower accuracy (Kim et al., 2014; Qin et al., 2014; Kim et al., 2017; Hulse-Kemp et al., 2018; Lee et al., 2022; Liao et al., 2022). High-fidelity (HiFi) long-read sequencing has substantially improved genome assembly accuracy, continuity, and completeness, enabling high-quality chromosome-scale assemblies across diverse pepper species (Shirasawa et al., 2023; Liu et al., 2023; Delorean et al., 2023; Chen et al., 2024; Zhang et al., 2025). In parallel, recent advances in pangenomics provide a powerful framework for characterizing genetic diversity within and across species. By integrating multiple high-quality genome assemblies, pangenomes capture a more complete repertoire of genetic variation than a single reference genome, thereby providing deeper insights into evolutionary dynamics, population diversity, and functional variation (Wang et al., 2023; Schreiber et al., 2024). Furthermore, graph-based pangenomes‒which represent multiple genome sequences within an interconnected graph‒have been shown to substantially improve the detection and genotyping of variants at the population scale, particularly large structural variants (SVs), compared to conventional linear reference approaches. These advances have facilitated trait discovery and genomics-assisted breeding in various crops (Zhou et al., 2022; Wang et al., 2023; Schreiber et al., 2024; Sun et al., 2026; Zhao et al., 2026).

Here, we present a *Capsicum* super-pangenome constructed from high-quality genome assemblies of 48 accessions, including 38 newly generated genomes, spanning all five domesticated taxa and their wild relatives. Integrating this genomic resource with population-scale genomic variation and extensive phenotypic characterization of 130 agronomic and metabolic traits‒including morphology and growth, carotenoids, capsaicinoids, and fruit volatiles‒provides insights into the genetic and functional diversity of the *Capsicum* genus. Collectively, our results illuminate the genomic basis of pepper domestication and metabolic diversification and establish valuable resources for genomics-assisted pepper improvement.

## Results

### Genome assembly and annotation of *Capsicum* accessions

To comprehensively capture genomic diversity across the *Capsicum* genus, we selected 38 representative accessions, primarily based on phylogenetic relationships among 500 core accessions (Liu et al., 2023). These comprised 23 cultivated accessions (five *C. pubescens*, four *C. baccatum* var. *pendulum*, five *C. frutescens*, four *C. chinense*, and five *C. annuum* var. *annuum*) and 15 wild accessions (one *C. rhomboideum*, four *C. chacoense*, five *C. baccatum* var. *baccatum*, one *C. galapagoense*, and four *C. annuum* var. *glabriusculum*). For each accession, we generated an average of 93.2 Gb (∼29×) of PacBio HiFi sequence data and 265.0 Gb (∼83×) of high-throughput chromosome conformation capture (Hi-C) data. Using these datasets, we generated chromosome-scale assemblies with an average contig N50 of 173.4 Mb (**Supplementary Fig. 1** and **Supplementary Table 1**). Assembly sizes ranged from 3.03 to 3.97 Gb, except for *C. rhomboideum*, which had a substantially smaller genome (∼1.7 Gb). Genome size was positively correlated with transposable element (TE) content (Pearson’s *r* = 0.63, *P* = 1.1 × 10^-6^). Quality assessments using Merqury (Rhie et al., 2020), BUSCO (Simão et al., 2015), and LAI (Ou et al., 2018) confirmed the high accuracy and completeness of the assemblies (**Supplementary Table 1**). We integrated the 38 newly generated assemblies with ten previously published high-quality genomes (Lee et al., 2022; Liao et al., 2022; Shirasawa et al., 2023; Liu et al., 2023; Delorean et al., 2023; Chen et al., 2024), yielding a dataset of 48 *Capsicum* genomes spanning diverse genetic groups: two *C. rhomboideum*, six *C. pubescens*, four *C. chacoense*, five *C. baccatum* var. *baccatum*, five *C. baccatum* var. *pendulum*, one *C. galapagoense*, five *C. frutescens*, four *C. chinense*, four *C. annuum* var. *glabriusculum*, and 12 *C. annuum* var. *annuum* accessions (**Fig. 1** and **Supplementary Table 1**). Genome-wide synteny was largely conserved across the 48 accessions, despite several large-scale rearrangements among lineages (**Fig. 1**).

**Fig. 1.**
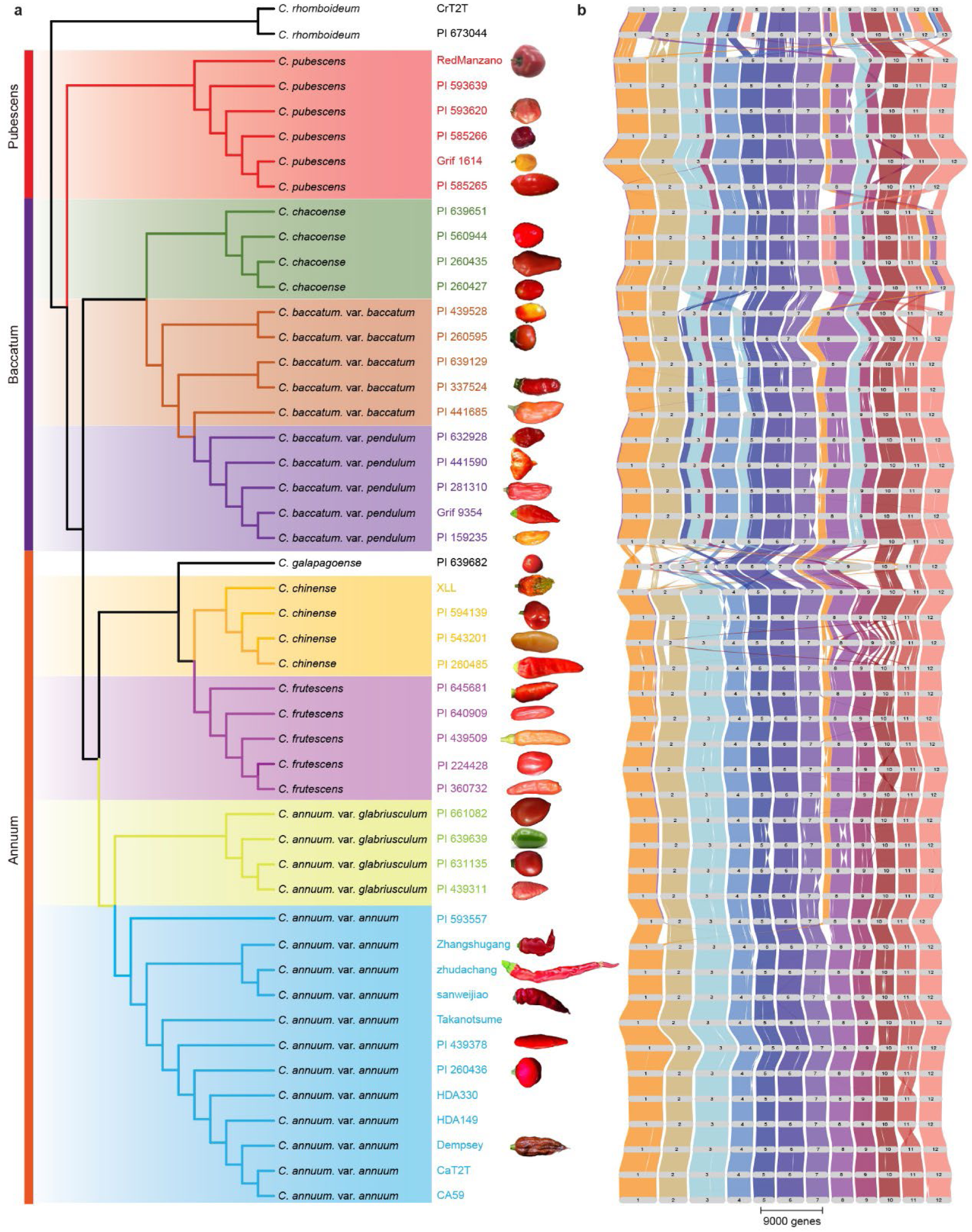
Phylogeny and genome synteny across the *Capsicum* genus. **a,** Phylogenetic relationships among the 48 *Capsicum* accessions included in this study. Fruit images are shown alongside the corresponding accessions where available. **b,** Genome-wide synteny among the 48 *Capsicum* genomes, generated using GENESPACE (https://github.com/jtlovell/GENESPACE). Horizontal gray bars represent chromosomes, and ribbons connecting adjacent genomes indicate syntenic relationships.

Gene prediction integrating *ab initio* prediction, transcript evidence, and protein homology identified an average of 37,005 protein-coding genes per genome. BUSCO analysis indicated that, on average, 95.74% of conserved single-copy plant orthologs were identified as complete, indicating high completeness of the gene predictions. For the ten previously published genomes, gene models were updated using Liftoff (Shumate and Salzberg, 2021), achieving annotation completeness comparable to that of the newly assembled genomes (**Supplementary Table 1**).

### Gene-based super-pangenome and pan-NLRome

To characterize gene-content diversity across the *Capsicum* genus, we constructed a gene-based super-pangenome from the 48 genomes and identified 72,687 gene families (pangenes). The pangene accumulation curve increased rapidly with the addition of the first several genomes and approached saturation at approximately 35 accessions (**Fig. 2a**). Based on their occurrence frequencies across the 48 genomes, pangenes were classified into four categories: core (10,427; 14.3%), soft-core (3,025; 4.2%), shell (55,952; 77.0%), and private (3,283; 4.5%) (**Fig. 2b**). The predominance of shell pangenes indicates extensive gene presence-absence variation (PAV) across *Capsicum*. Functional enrichment analyses revealed that core pangenes were primarily involved in essential cellular and metabolic processes, whereas soft-core, shell, and private pangenes were enriched for functions related to environmental adaptation and stress responses (**Supplementary Fig. 2**).

**Fig. 2.**
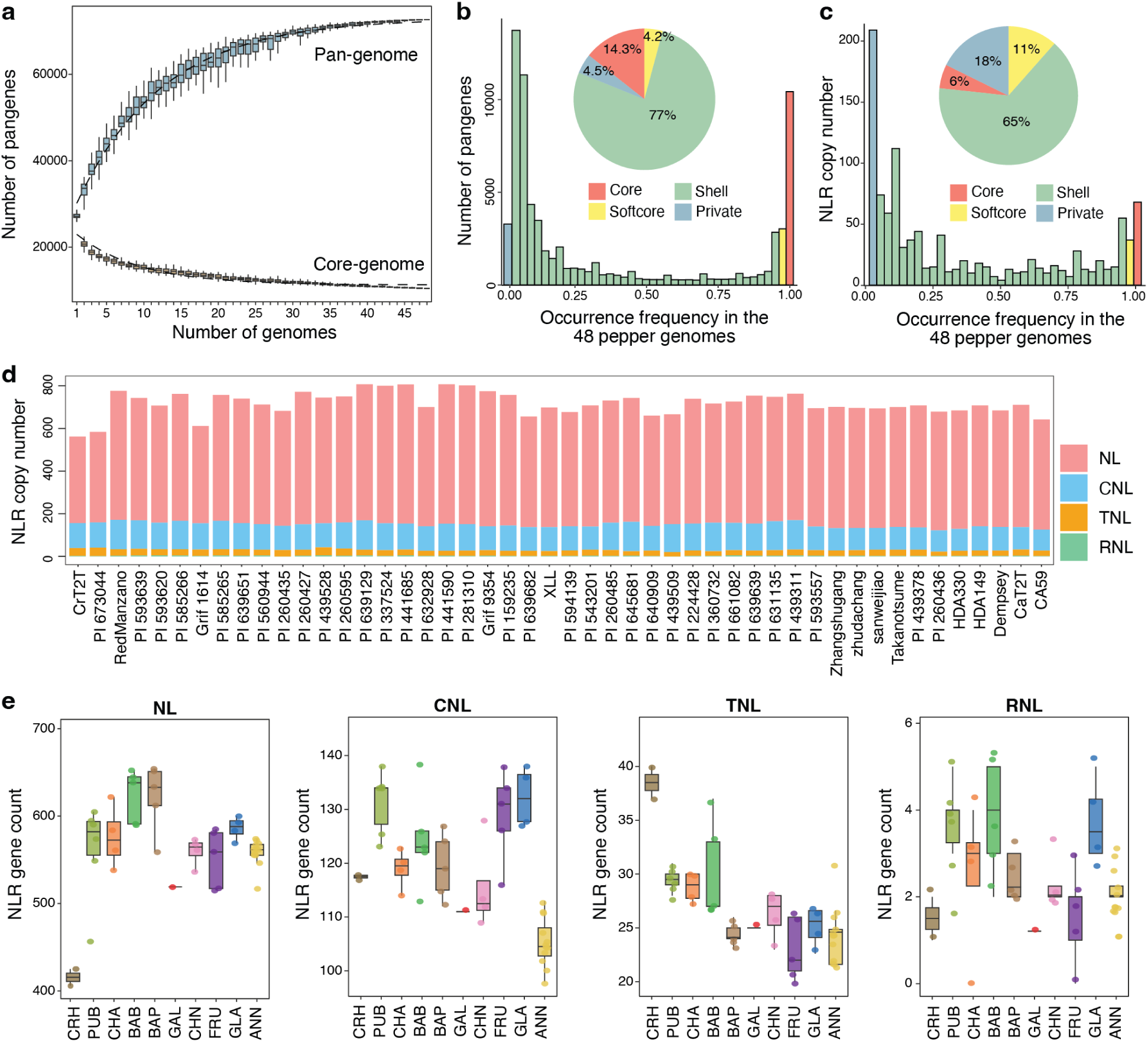
Gene-based super-pangenome and pan-NLRome of the *Capsicum* genus. **a,** Pangenome expansion and core-genome contraction with increasing numbers of genomes. For each given number of genomes, 100 random combinations of accessions were sampled. Boxplots show the distributions across random combinations, and dashed lines indicate the fitted accumulation and contraction curves for the pangenome and core genome, respectively. **b,c,** Distribution of pangene (**b**) and NLR orthogroup (**c**) occurrence frequencies across the 48 genomes. Pangenes and NLR orthogroups were classified as core (present in all 48 genomes), soft-core (present in 43–47 genomes), shell (present in 2–42 genomes), and private (present in a single genome). Insets show the proportions of pangenes and NLR orthogroups in the four categories. **d,** NLR copy numbers across the 48 *Capsicum* accessions, with NLRs classified into NL, CNL, TNL, and RNL subclasses. **e,** Comparison of NLR copy numbers among major *Capsicum* groups. For each boxplot, the lower and upper bounds indicate the first and third quartiles, respectively, the centerline indicates the median and the whiskers extend to 1.5× interquartile range. CRH, *C. rhomboideum*; PUB, *C. pubescens*; CHA, *C. chacoense*; BAB, *C. baccatum* var. *baccatum*; BAP, *C. baccatum* var. *pendulum*; GAL, *C. galapagoense*; CHN, *C. chinense*; FRU, *C. frutescens*; GLA, *C. annuum* var. *glabriusculum*; ANN, *C. annuum* var. *annuum*.

We reconstructed a time-calibrated phylogeny of *Capsicum*, together with tomato, potato, and eggplant, using 5,020 single-copy orthologs. The phylogeny placed *C. rhomboideum* as the earliest-diverging lineage within *Capsicum*, followed by *C. pubescens*, with the remaining species forming a more recently diversified clade. Divergence-time estimates indicated that *C. rhomboideum* diverged from the remaining *Capsicum* lineages approximately 13.56 million years ago (Ma), followed by *C. pubescens* at approximately 3.90 Ma, whereas the remaining *Capsicum* lineages diversified approximately 1.24-2.42 Ma (**Supplementary Fig. 3**). Within this phylogenetic framework, comparison of pangene distributions across major *Capsicum* lineages revealed that, although a substantial proportion of pangenes were shared among lineages, each lineage also contained a distinct set of lineage-specific pangenes (**Supplementary Fig. 4**).

To investigate disease-resistance gene diversity, we constructed a pan-NLRome from the 48 *Capsicum* genomes. In total, 6,533 NLR genes were identified and clustered into 1,115 orthogroups (pangenes). Most NLR pangenes were classified as shell or private (**Fig. 2c**), revealing extensive variation in NLR repertoires among accessions. The pan-NLRome accumulation curve approached saturation at approximately 40 accessions (**Supplementary Fig. 5**). NLR genes were further classified into four subclasses: TIR-NLR (TNL), CC-NLR (CNL), CCR-NLR (RNL), and NB-and-LRR-only (NL). Across all accessions, NLs represented the predominant subclass, followed by CNLs, whereas TNLs and RNLs were comparatively rare (**Fig. 2d**). Comparison of NLR repertories across major *Capsicum* groups revealed broadly similar distributions of NLR subclasses but substantial lineage-specific variation in NLR copy number (**Fig. 2e**). *C. pubescens*, *C. baccatum* var. *baccatum*, and *C. baccatum* var. *pendulum* contained relatively large NL repertoires, whereas *C. rhomboideum* possessed the fewest NL genes. CNLs were more abundant in *C. pubescens*, *C. frutescens*, and *C. annuum* var. *glabriusculum* than in the other groups. TNLs were most abundant in *C. rhomboideum* and *C. pubescens* but less abundant in several more recently diverged lineages, including *C. baccatum* var. *pendulum*, *C. galapagoense*, *C. frutescens*, and *C. annuum* var. *annuum*. RNLs were consistently the least abundant subclass across all groups. We further compared NLR repertoires along two independent domestication trajectories represented in our dataset: *C. baccatum* var. *baccatum* versus domesticated *C. baccatum* var. *pendulum*, and *C. annuum* var. *glabriusculum* versus domesticated *C. annuum* var. *annuum*. In both comparisons, domesticated accessions generally contained fewer NLR genes than their corresponding wild relatives (**Fig. 2e**), suggesting that domestication was accompanied by a reduction in NLR copy number.

Together, the gene-based super-pangenome and pan-NLRome provide a framework for characterizing gene-content and immune-receptor diversity across *Capsicum*, facilitating comparative analyses of genome evolution and providing a valuable resource for the genetic improvement of agronomically important traits, particularly disease resistance.

### Graph-based super-pangenome construction and SV genotyping

To comprehensively capture structural variation across *Capsicum*, we constructed a graph-based super-pangenome using 46 high-quality genomes, excluding the two highly divergent *C. rhomboideum* accessions. The graph contained a large set of SVs (≥20 bp), including 2,012,901 insertions and 1,498,594 deletions (**Supplementary Table 2**). SV abundance varied substantially among accessions, ranging from 34,033 to 273,156 variants per accession (**Supplementary Table 3**), highlighting extensive structural diversity across *Capsicum*. Functional annotation revealed that 90.8% of SVs were located in intergenic regions, whereas a subset was predicted to have potentially disruptive effects, including 6,914 frameshift, 4,045 stop-gained, and 14,587 exon-loss variants (**Supplementary Table 4**).

To characterize SV dynamics at the population level, we genotyped these SVs across 500 core *Capsicum* accessions using short-read resequencing data (Liu et al., 2023). Given the challenges of short-read SV genotyping, particularly in repetitive and structurally complex regions, we evaluated genotyping performance using 35 accessions for which both chromosome-scale assemblies and short-read resequencing data were available. Using assembly-derived SV genotypes as a benchmark, 961,086 insertions and 772,159 deletions achieved F1 scores >0.7 (**Supplementary Table 2**) and were retained as a high-confidence SV set. Allele frequencies estimated from short-read genotyping for these SVs were highly concordant with those derived from genome assemblies (**Fig. 3a,b**). Furthermore, phylogenetic relationships among the 500 core accessions inferred from these high-confidence SVs (**Supplementary Fig. 6**) were largely consistent with those inferred from genome-wide SNPs (Liu et al., 2023). We therefore used this high-confidence SV set for subsequent population-level analyses.

**Fig. 3.**
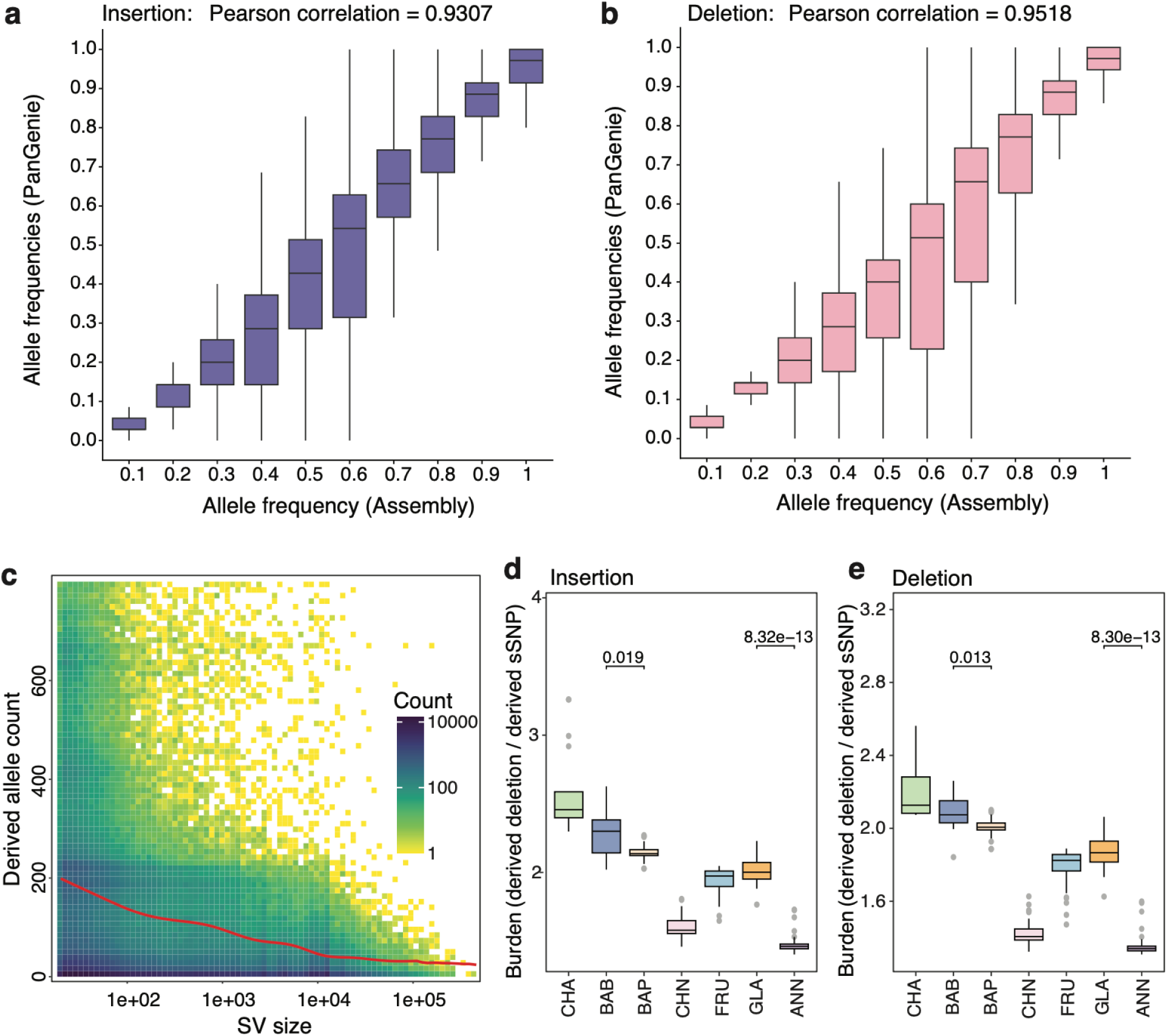
Landscape and evolutionary patterns of structural variation in the *Capsicum* super-pangenome. **a,b,** Concordance of allele frequencies between assembly-derived SV calls and graph-based SV genotypes inferred from short-read sequencing for insertions (**a**) and deletions (**b**). **c,** Relationship between SV size and derived allele count. Colors indicate the number of SVs in each bin on a logarithmic scale. The red curve represents a fitted generalized additive model. **d,e,** Insertion (**d**) and deletion (**e**) burdens across major *Capsicum* groups. Numbers above brackets indicate P values for comparisons between wild and domesticated groups (two-sided Student’s *t*-tests). For each boxplot, the lower and upper bounds indicate the first and third quartiles, respectively, the centerline indicates the median and the whiskers extend to 1.5× interquartile range. PUB, *C. pubescens*; CHA, *C. chacoense*; BAB, *C. baccatum* var. *baccatum*; BAP, *C. baccatum* var. *pendulum*; FRU, *C. frutescens*; CHN, *C. chinense*; GLA, *C. annuum* var. *glabriusculum*; ANN, *C. annuum* var. *annuum*.

### Evolutionary signatures and domestication-related patterns of SVs

Population-scale SV genotyping revealed substantial variation in SV abundance among *Capsicum* species and breeding groups (**Supplementary Fig. 7** and **Supplementary Table 5**). To investigate patterns of SV differentiation during pepper domestication, we compared SV allele frequencies along two independent domestication trajectories: *C. baccatum* var. *baccatum* to *C. baccatum* var. *pendulum*, and *C. annuum* var. *glabriusculum* to *C. annuum* var. *annuum*. Using a false discovery rate threshold of <0.001 and a >twofold difference in allele frequency, we identified 27,035 and 37,908 differentiated SVs in these two comparisons, respectively, revealing extensive shifts in SV frequencies between wild and domesticated groups. These differentiated SVs overlapped 4,106 and 5,648 genes, respectively (**Supplementary Table 6**). Notably, genes associated with differentiated SVs in both domestication trajectories included *ACS1* (*Caa02_01g01760*) and *CCR* (*Caa02_01g08820*). *ACS1* encodes 4-coumarate:CoA ligase (4CL), which catalyzes the formation of 4-coumaroyl-CoA, a key intermediate in the phenylpropanoid branch of capsaicinoid biosynthesis (Vergnano et al., 2026). *CCR* encodes cinnamoyl-CoA reductase, an enzyme involved in phenylpropanoid metabolism and the production of capsaicinoid precursors (Reddy et al., 2014). The occurrence of differentiated SVs associated with these genes in both domestication trajectories highlights recurrent changes involving the phenylpropanoid pathway and suggests a potential link between SV differentiation and capsaicinoid metabolism during pepper domestication.

We next investigated the evolutionary dynamics and potential fitness consequences of SVs by comparing derived allele frequency patterns between SNPs and SVs. We found a greater enrichment of rare derived alleles among SVs than among SNPs (**Supplementary Fig. 8a**), a pattern consistent with stronger purifying selection against SVs (Collins and Talkowski, 2025). Derived allele count also decreased with increasing SV size (**Fig. 3c**), indicating stronger evolutionary constraint on larger SVs, which may reflect their greater potential to disrupt genome structure and function.

We then quantified SV burden across major *Capsicum* populations as the number of derived insertion and deletion alleles carried by each accession, normalized by the number of derived synonymous SNP alleles. Both insertion and deletion burdens varied substantially among populations (**Fig. 3d,e**). *C. chacoense* exhibited the highest burdens, followed by *C. baccatum* var. *baccatum* and *C. baccatum* var. *pendulum*. *C. frutescens* and *C. annuum* var. *glabriusculum* displayed intermediate burdens, whereas *C. chinense* and *C. annuum* var. *annuum* exhibited the lowest burdens. Along the two domestication trajectories represented in our dataset, domesticated populations carried significantly lower SV burdens than their corresponding wild progenitors (**Fig. 3d,e**). In contrast, nonsynonymous SNP (nSNPs) burdens were higher in domesticated populations than in their wild counterparts (**Supplementary Fig. 8b**). These contrasting patterns of reduced SV burden and elevated nSNP burden in domesticated populations may reflect the effects of demographic history and differences in selection on these two classes of genetic variation during domestication and subsequent breeding.

### Graph-based super-pangenome empowers association mapping of complex traits

We generated a comprehensive phenotypic dataset comprising 130 traits across the pepper core diversity panel, including 25 morphological and growth traits and 105 specialized metabolites— four capsaicinoids, eight carotenoids, and 93 fruit volatile compounds (**Fig. 4a** and **Supplementary Tables 7-10**). These data revealed extensive phenotypic diversity across the germplasm and pronounced differences between wild and domesticated accessions. For instance, fruit shape differed markedly between domesticated accessions and their wild relatives (**Fig. 4b**), reflecting substantial morphological diversification during domestication. Pedicel orientation also differed, with horizontal and downward pedicels occurring at substantially higher frequencies in domesticated accessions than in their wild progenitors (**Fig. 4c**).

**Fig. 4.**
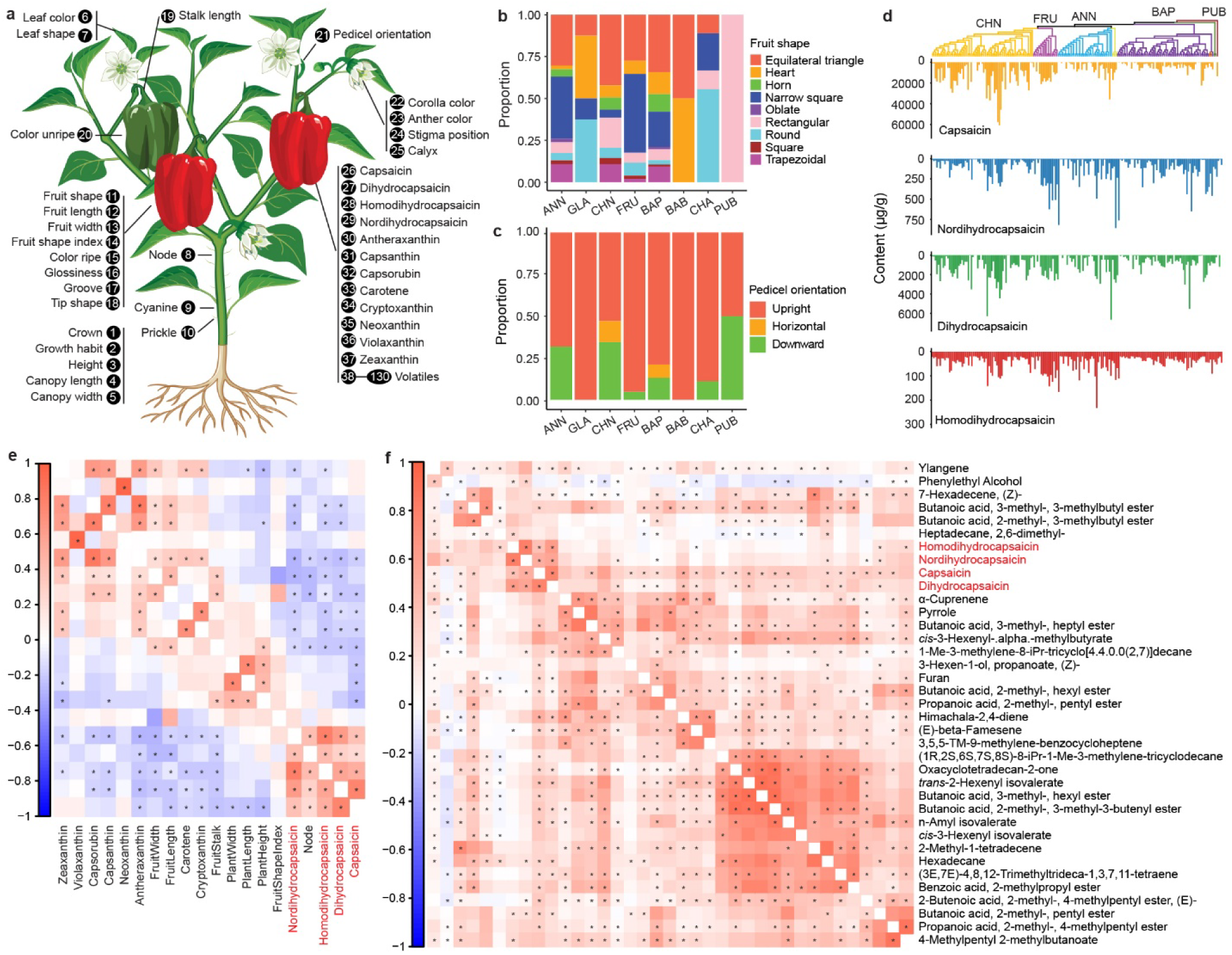
A comprehensive phenome of the pepper core collection. **a**, Overview of the phenotypic traits measured in this study. **b,c**, Distribution of fruit shape (**b**) and pedicel orientation (**c**) across *Capsicum* species and varieties. **d**, Contents of the four major capsaicinoids across the pepper core collection, with accessions arranged according to their phylogenetic relationships and taxonomic groups. **e**, Correlations between capsaicinoid levels, major agronomic traits, and carotenoid contents. **f**, Correlations between capsaicinoid levels and fruit volatile compounds. Colors in **e** and **f** indicate Pearson correlation coefficients, and asterisks indicate significant correlations (*P* < 0.05). ANN, *C. annuum* var. *annuum*; GLA, *C. annuum* var. *glabriusculum*; CHN, *C. chinense*; FRU, *C. frutescens*; BAP, *C. baccatum* var. *pendulum*; BAB, *C. baccatum* var. *baccatum*; CHA, *C. chacoense*; PUB, *C. pubescens*.

The four major capsaicinoids—capsaicin, dihydrocapsaicin, homodihydrocapsaicin, and nordihydrocapsaicin—varied widely across the genus, underscoring extensive diversity in pepper pungency (**Fig. 4d**). Capsaicin, a major contributor to pungency, exhibited the greatest variation and accumulated to the highest levels in *C. chinense*. In contrast, the other three major capsaicinoids accumulated to the highest levels in *C. annuum* var. *annuum*, revealing substantial interspecific variation in capsaicinoid composition. Predominantly negative correlations were observed between capsaicinoid levels and both carotenoid contents and several agronomic traits (**Fig. 4e**). The four major capsaicinoids were also positively correlated (*P* < 0.05, *r* ≥ 0.2) with 33 volatile compounds spanning diverse chemical classes, including terpenoids, branched-chain esters, and C6 green-leaf volatiles (**Fig. 4f**).

Integrating this phenotypic dataset with population-scale SNPs and SVs enabled comprehensive genome-wide association studies (GWAS) across diverse agronomic and metabolic traits. In total, we identified 2,831 significant associations across 107 of the 130 traits evaluated (**Supplementary Table 11**). Of these, 56 traits showed significant associations with SNPs only, whereas 51 exhibited significant associations with both SNPs and SVs, demonstrating that structural variation provides complementary information for dissecting the genetic architecture of agronomic and metabolic diversity in *Capsicum*.

Among the associated loci, a prominent signal on chromosome 4 (peak position: Chr04:31,627,338; LD interval: 31.48–32.63 Mb) was associated with capsorubin content (**Supplementary Fig. 9a**). The alternative allele, detected exclusively in *C. chinense*, was associated with significantly higher capsorubin levels than the reference allele present in *C. chinense* and other species, including *C. baccatum* var. *pendulum* and *C. frutescens* (**Supplementary Fig. 9b,c**). Among the 21 annotated genes within the LD interval, *Caa02_04g09950*, encoding an ethylene-responsive transcription factor RAP2-13-like protein (hereafter RAP2), emerged as a strong candidate based on its substantially higher expression in fruit than neighboring genes (**Supplementary Fig. 9d**). Functional validation using sustained virus-induced gene silencing (VIGS) showed that silencing *RAP2* resulted in an orange-red fruit phenotype and markedly reduced *RAP2* transcript abundance relative to red-ripe control fruits (**Supplementary Fig. 9e,f**). Targeted metabolomic profiling further revealed pronounced reductions in the major red carotenoids capsanthin and capsorubin, together with decreases in several esterified carotenoids, including zeaxanthin dipalmitate and violaxanthin dimyristate. Conversely, several upstream carotenoid intermediates, including antheraxanthin, neoxanthin, and lutein, accumulated to significantly higher levels following *RAP2* silencing (**Supplementary Table 12**). Together, these results support a role for *RAP2* in regulating carotenoid metabolism and promoting the accumulation of downstream red carotenoids in pepper fruit.

For capsaicinoids, a significant locus spanning 25.50–27.26 Mb on chromosome 11 was associated with homodihydrocapsaicin content. This interval harbors four tandemly duplicated enoyl-[acyl-carrier-protein] reductase (ENR) genes (*Caa02_11g06430–Caa02_11g06460*), which encode key enzymes involved in the fatty acid biosynthetic pathway that supplies precursors for capsaicinoid side-chain formation (**Supplementary Fig. 10a**). Accessions carrying the alternative allele of the lead SNP accumulated significantly higher levels of all four major capsaicinoids than those carrying the reference allele (**Supplementary Fig. 10b**), suggesting that genetic variation at this locus may contribute broadly to capsaicinoid accumulation.

Integration of SNP- and SV-based GWAS further resolved a structurally complex locus associated with the sesquiterpene ylangene on chromosome 9 (∼13–16 Mb). Whereas SNP-based mapping identified a broad high-LD region, SV-based mapping revealed a local breakdown of LD within this interval (**Fig. 5a**), highlighting its structural complexity. Comparative genomic analysis revealed extensive copy-number variation (CNV) within a terpene synthase (TPS) gene cluster, with tandem amplification occurring after the divergence of *C. rhomboideum* and *C. pubescens*, resulting in a marked expansion of TPS copy number in the remaining *Capsicum* lineages (**Fig. 5b**). Synteny analysis further showed that the homologous genomic segment is located on chromosome 3 in several early-diverging lineages, including *C. rhomboideum*, *C. pubescens*, *C. chacoense*, and *C. baccatum*, but on chromosome 9 in the Annuum clade, including *C. galapagoense*, *C. frutescens*, *C. chinense*, and *C. annuum*, consistent with an interchromosomal translocation. Moreover, a lineage-specific inversion within this interval was identified in a subset of *C. annuum* accessions. Accessions carrying the alternative alleles of both the peak SNP and lead SV accumulated significantly higher levels of ylangene (**Fig. 5c**). Multiple potentially functional variants, including missense, splice-site, and stop-gain variants, were identified within genes of the TPS cluster. Expression profiling further prioritized *Caa02_09g05220* as the only TPS gene consistently expressed throughout fruit development. Notably, *Caa02_09g05220* harbored 20 missense and five splice-site variants (**Fig. 5d**). Together, these results reveal a highly dynamic TPS locus shaped by copy-number expansion, large-scale structural rearrangements, and extensive sequence variation, which may have collectively contributed to the diversification of sesquiterpene profiles across *Capsicum*.

**Fig. 5.**
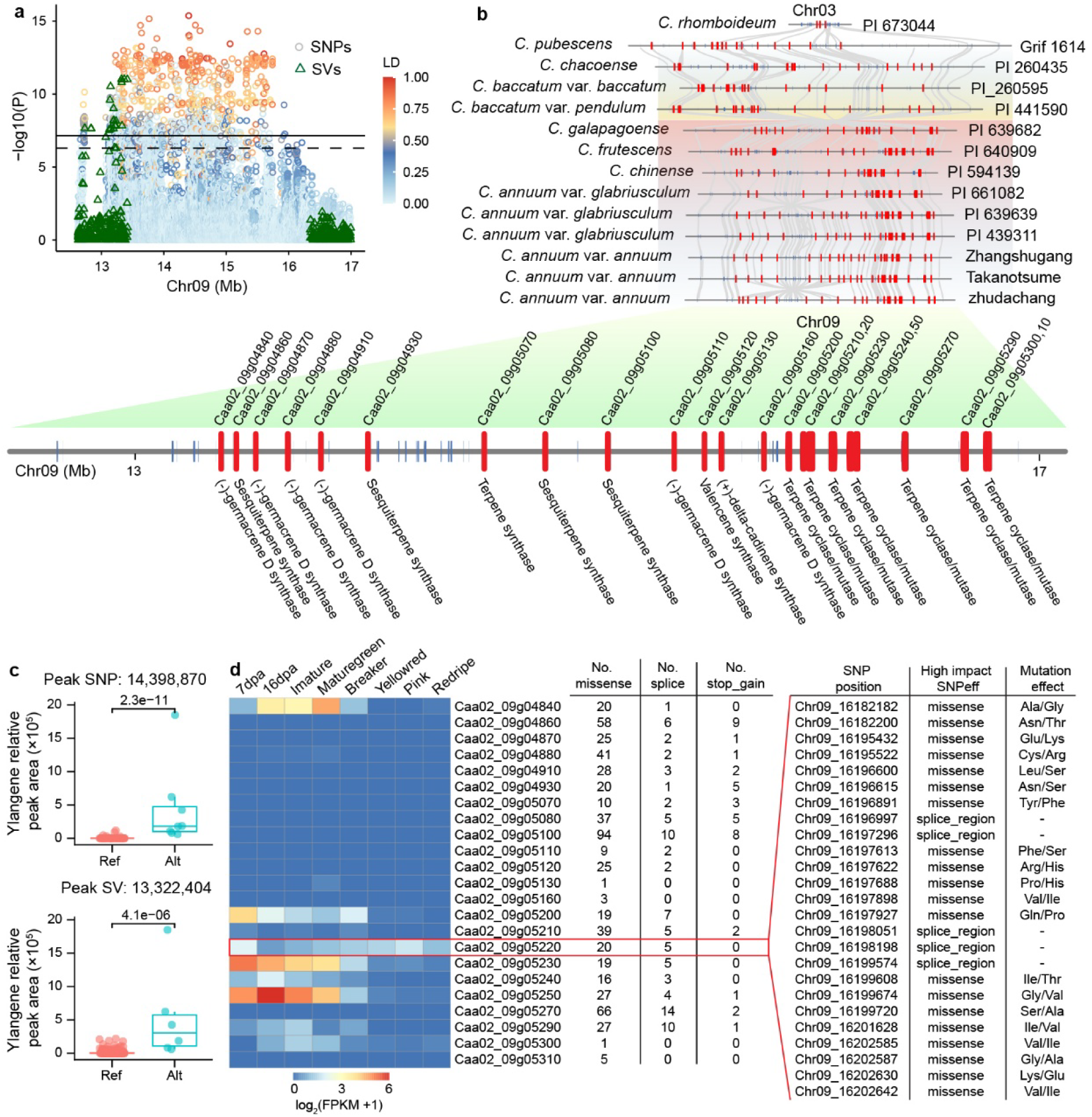
Evolutionary and functional dissection of a terpene synthase gene cluster associated with ylangene accumulation. **a**, Regional Manhattan plot showing SNP-based (circles) and SV-based (triangles) GWAS signals for ylangene content. Circle colors indicate linkage disequilibrium (LD; *r*²) between the lead SNP (Chr09:14,398,870) and neighboring SNPs. **b**, Synteny of the associated locus across representative *Capsicum* genomes. Terpene synthase (TPS) genes are highlighted in red. **c**, Ylangene levels in accessions carrying the reference (Ref) or alternative (Alt) alleles of the lead SNP (Chr09:14,398,870; top) and lead SV (Chr09:13,322,404; bottom). *P* values were calculated using two-sided Wilcoxon rank-sum tests. For each boxplot, the lower and upper bounds indicate the first and third quartiles, respectively, the centerline indicates the median and the whiskers extend to 1.5× interquartile range. **d**, Expression profiles of TPS genes within the associated interval during fruit development, together with predicted potentially high-impact variants, including missense, splice-site and stop-gained variants.

### Pangenomic dissection of capsaicinoid biosynthetic genes

To dissect the genetic architecture underlying capsaicinoid biosynthesis, we systematically annotated potential capsaicinoid biosynthetic genes (CBGs) across the 48 *Capsicum* genomes. We identified 148 CBG families, substantially expanding the repertoire represented in individual genomes (∼97 per genome; range, 86-106) and revealing extensive PAV and CNV across the pathway (**Supplementary Table 13**). Of these families, 54 (36.5%) were classified as core and 27 (18.2%) as soft-core, substantially higher than the corresponding genome-wide proportions of 14.3% and 4.2%, respectively, indicating that the gene repertoire underlying capsaicinoid biosynthesis is more broadly conserved across *Capsicum* lineages than the genome-wide gene repertoire.

We next examined CBG expression in the placenta, the primary site of capsaicinoid biosynthesis, at different stages of fruit development in the pungent cultivar “Zhangshugang” and two non-pungent mutants, c68 and ss97. We identified 15 CBGs that were consistently upregulated in the pungent cultivar relative to both non-pungent mutants across all developmental stages (**Fig. 6a**), representing a 28.3-fold enrichment of CBGs relative to the genome-wide background (Fisher’s exact test, *P* = 7.4 × 10^-18^). Notably, nine of these 15 CBGs were tandemly duplicated, representing a 3.0-fold enrichment of tandem duplicates relative to the genome-wide background (Fisher’s exact test, *P* = 8.7 × 10^-4^) (**Supplementary Table 14**). These tandemly duplicated genes participate in multiple key steps of the capsaicinoid biosynthetic pathway (**Fig. 6b**), highlighting the contribution of tandem duplication to the expansion and diversification of the genetic repertoire underlying pepper pungency.

**Fig. 6.**
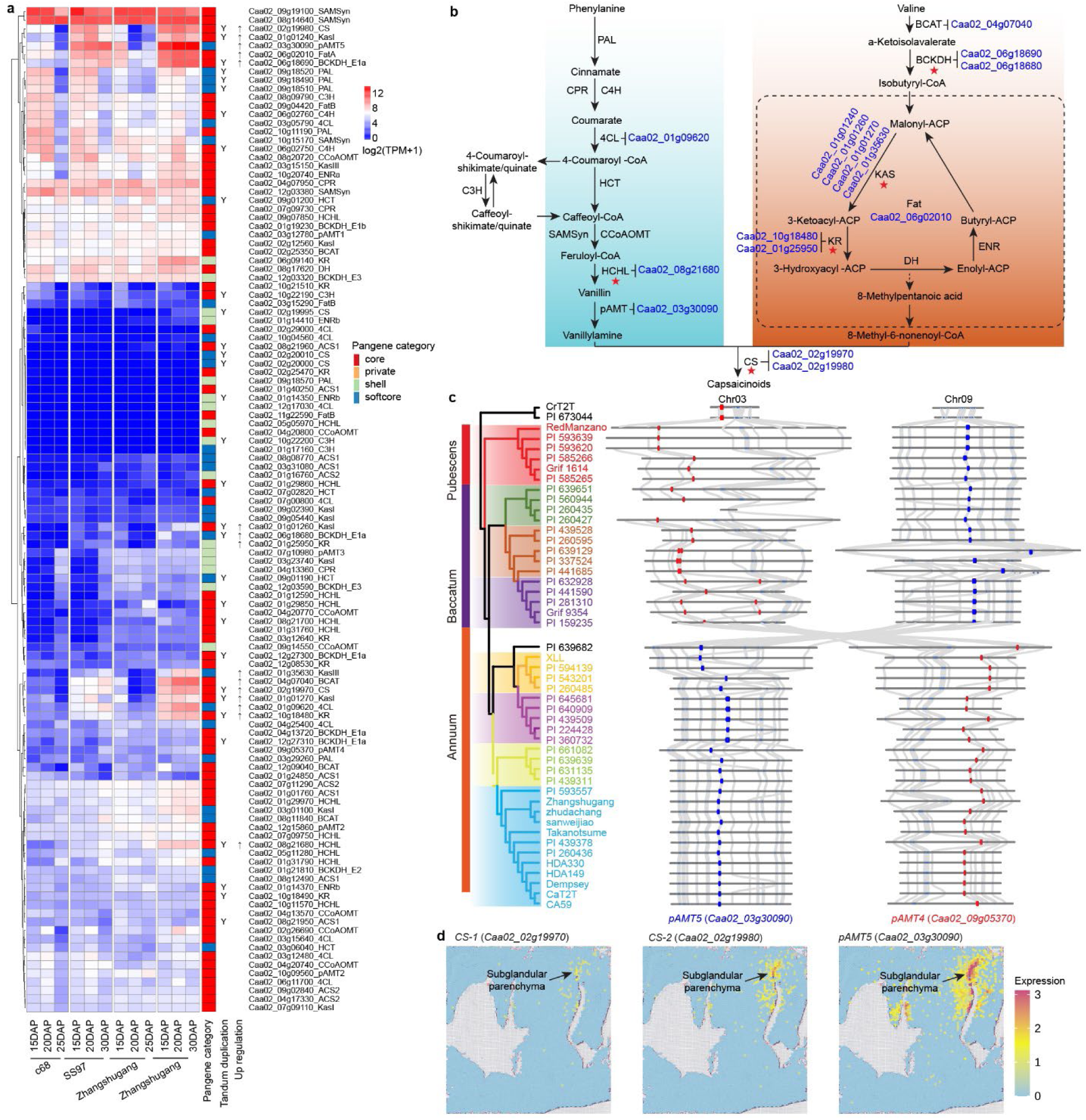
Expression and evolution of capsaicinoid biosynthetic genes. **a**, Expression heatmap of capsaicinoid biosynthetic genes (CBGs) in the placenta of the pungent cultivar ‘Zhangshugang’ and two non-pungent mutants, c68 and ss97. Expression values are shown as log_2_(TPM + 1). “Y” denotes tandemly duplicated genes, and “↑” indicates CBGs consistently upregulated in ‘Zhangshugang’ relative to both non-pungent mutants. **b**, Capsaicinoid biosynthetic pathway. Pathway steps involving tandemly duplicated genes are marked with red stars, and genes preferentially upregulated in the pungent cultivar are highlighted in blue. **c**, Regional synteny surrounding *pAMT4* (red) and *pAMT5* (blue) across *Capsicum* genomes, illustrating an interchromosomal rearrangement between chromosomes 3 and 9. **d**, Spatial expression patterns of *CS-1*, *CS-2* and *pAMT5* in pepper fruit at 25 days after pollination (DAP).

Among these nine tandemly duplicated CBGs were two capsaicin synthase (CS) genes, *Caa02_02g19970* and *Caa02_02g19980* (hereafter *CS-1* and *CS-2*), located within the *Pun1* locus. *CS-*2 corresponds to *AT3*, previously shown to underlie the *Pun1* locus (Stewart et al., 2005). Comparative analysis across the 48 genomes identified six *AT3* paralogous lineages at the *Pun1* locus (**Supplementary Table 15** and **Supplementary Fig. 11**). These lineages showed contrasting evolutionary patterns, ranging from widespread retention to partial loss or pseudogenization across *Capsicum*. *CS-1* and *CS-2* belonged to the same *AT3* lineage, but their orthologues formed two distinct phylogenetic clades (**Supplementary Fig. 12**), indicating that the duplication giving rise to this gene pair predated the diversification of the sampled *Capsicum* lineages. The two genomes of the non-pungent species *C. rhomboideum* lacked intact copies of both *CS-1* and *CS-2*, retaining only a single pseudogenized copy. In contrast, 42 of the remaining 46 genomes retained intact copies of both genes, whereas the other four carried partial or disruptive variants affecting one of the two copies. Other *AT3* lineages showed more extensive loss or pseudogenization. Expression patterns further distinguished the *CS-1*/*CS-2* lineage from the other *AT3* paralogous lineages. Among the four lineages retaining intact genes at the *Pun1* locus in ‘Zhudachang’, only *CS-1* and *CS-2* were strongly expressed during capsaicinoid accumulation (**Supplementary Fig. 13**). Moreover, *CS-2* was consistently expressed at higher levels than *CS-1* in the placenta of the pungent cultivar ‘Zhangshugang’, indicating expression divergence between the duplicated copies. Together, these patterns reveal extensive evolutionary remodeling of the *Pun1* locus through gene duplication, differential retention and loss, pseudogenization, and expression divergence among *AT3* paralogous lineages, with *CS-1* and, particularly, *CS-2* emerging as the predominant copies associated with capsaicinoid biosynthesis.

In addition to *CS*, aminotransferase (*pAMT*) is another key determinant of pepper pungency (Tsurumaki and Sasanuma, 2019). Among the nine *pAMT* copies identified in the *Capsicum* pangenome (**Supplementary Table 13**), *pAMT5* (*Caa02_03g30090*) was the only member preferentially upregulated in pungent fruit. Comparative genomic analysis revealed that *pAMT5* was absent from the non-pungent *C. rhomboideum*. Moreover, an interchromosomal translocation between chromosomes 3 and 9 that distinguishes the Baccatum and Annuum clades reorganized the genomic context surrounding *pAMT4* and *pAMT5* (**Fig. 6c**), suggesting a potential relationship between large-scale chromosomal rearrangement and regulatory divergence at this locus.

Spatial transcriptomic analysis using a public dataset (Han et al., 2026) further showed that *CS-1*, *CS-2*, and *pAMT5* were specifically enriched in the subglandular parenchyma, a distinct cell layer within the placental septum (**Fig. 6d**), revealing a spatially localized transcriptional program associated with capsaicinoid biosynthesis. Together, these findings indicate that the broadly conserved gene repertoire underlying capsaicinoid biosynthesis has been shaped by gene duplication, structural rearrangement, and regulatory divergence, with key pathway genes exhibiting highly localized expression within the placenta.

### Pangenome-enabled genomic prediction of complex traits in pepper

We leveraged genetic variation captured by the pepper graph-based super-pangenome to develop genomic prediction models for 130 phenotypes spanning agronomic traits, capsaicinoids, carotenoids, and volatile organic compounds (VOCs). For each trait, informative genome-wide variants were selected based on feature importance, and prediction models were constructed using either SNP-only or combined SNP+SV marker panels. The resulting trait-specific panels comprised only several hundred markers (mean = 769; **Supplementary Table 16**). SVs constituted a small proportion of the selected markers (1.1-3.6%), consistent with their lower abundance relative to SNPs in the pangenome variation dataset. Five-fold cross-validation demonstrated robust predictive performance across the 130 traits using these compact marker panels. The SNP+SV models achieved a mean Pearson correlation of 0.78 between predicted and observed phenotypic values, ranging from 0.53 to 0.98 across traits (**Supplementary Table 16**). Incorporating SVs provided modest, trait-dependent improvements in prediction accuracy, with SNP+SV models outperforming SNP-only models for 36 of the 130 traits. Collectively, these results demonstrate that pangenome-derived genetic variation can support genomic prediction across diverse agronomic and metabolic traits in pepper. The predictive performance achieved with relatively compact marker panels indicates that informative variants can be identified from the pangenome variation dataset for a broad range of phenotypes. Although SVs improved prediction for a subset of traits, their overall contribution beyond SNPs was modest, suggesting that SNPs capture much of the predictive signal, with SVs providing complementary information for specific traits.

## Discussion

The *Capsicum* super-pangenome, integrating 48 high-quality genome assemblies spanning all five domesticated taxa and their wild relatives, establishes a comprehensive genomic framework for capturing diversity beyond that represented by a single reference genome. By coupling this resource with population-scale genomic variation and extensive phenotypic data, our study provides a multidimensional view of how gene-content variation, structural rearrangements, and regulatory divergence contribute to agronomic and metabolic diversity in pepper.

The pan-NLRome and population-scale SV analyses revealed that domestication has reshaped multiple layers of genomic diversity in *Capsicum*. Along two independent domestication trajectories, domesticated accessions contained fewer NLRs than their wild progenitors, consistent with similar reductions reported in cucumber (Zhang and Deng, 2026). SVs, meanwhile, were enriched for rare derived alleles, with derived allele frequencies decreasing with increasing SV size, consistent with stronger purifying selection against larger structural changes. Domesticated peppers carried lower SV burdens than their corresponding wild progenitors, a pattern also reported in cucumber (Zhao et al., 2026), suggesting that demographic processes and selection associated with domestication and subsequent breeding have contributed substantially to changes in the SV landscape.

Integrating SNPs and SVs in GWAS provided a more comprehensive view of the genetic architecture underlying complex agronomic and metabolic traits. Significant SV associations were detected for 51 traits, demonstrating that structural variation provides complementary information to SNP-based mapping. The structurally complex *TPS* locus associated with ylangene exemplifies this added resolution. Whereas SNP-based mapping identified a broad region of high linkage disequilibrium that obscured the underlying genetic architecture, SV-based analysis revealed a dynamic locus shaped by copy-number expansion, interchromosomal translocation, and lineage-specific inversion. In parallel, functional analysis of *RAP2*, identified from a locus associated with capsorubin content, revealed a role for this ethylene-responsive transcription factor in carotenoid metabolism and the accumulation of red carotenoids in pepper fruit. This finding expands the regulatory network underlying fruit pigmentation beyond canonical carotenoid biosynthetic enzymes.

Our pangenomic dissection of capsaicinoid biosynthesis further revealed extensive evolutionary remodeling of the pathway through gene duplication, differential retention and loss, structural rearrangement, and expression divergence. At the *Pun1* locus, contrasting evolutionary trajectories among *AT3* paralogous lineages, together with the broad retention and strong expression of *CS-1* and *CS-2*, illustrate how gene duplication and subsequent divergence have reshaped a key locus underlying pungency. The distinct expression pattern of *pAMT5*, together with large-scale rearrangement of its genomic context, further highlights the potential contributions of structural and regulatory variation to pathway evolution. Spatial transcriptomics further revealed that key pathway genes are specifically expressed in the subglandular parenchyma of the placental septum, revealing a highly localized transcriptional program associated with capsaicinoid biosynthesis. These findings refine the cellular context of specialized metabolism in pepper and extend previous models proposing that duplication and functional divergence of ancestral metabolic genes contributed to the evolution of capsaicinoid biosynthesis (Kim et al., 2014; Kusaka et al., 2024).

Beyond these evolutionary and functional insights, the super-pangenome provides a foundation for translating expanded genomic diversity into pepper improvement. The high genomic prediction accuracy achieved with relatively compact marker panels across diverse agronomic and metabolic traits highlights the potential of pangenome-derived variation to inform genomic selection. Collectively, our study establishes a comprehensive pangenomic framework that illuminates the genomic consequences of *Capsicum* domestication and the genetic basis of metabolic diversification while providing valuable resources for genomics-assisted improvement. Further integration of structural variation with functional genomic and evolutionary information should enhance the identification and deployment of genetic diversity to develop resilient, high-quality pepper varieties.

## Materials and Methods

### Genome sequencing and de novo assembly

High-molecular-weight (HMW) DNA was extracted from young, fresh leaves of 38 *Capsicum* accessions using the cetyltrimethylammonium bromide (CTAB) method (Doyle and Doyle, 1987). PacBio SMRT libraries were constructed from HMW DNA using the SMRTbell Express Template Prep Kit 2.0 (PacBio, USA) and sequenced on the PacBio Sequel II platform to generate HiFi reads. Hi-C libraries were prepared following the Proximo Hi-C Plant protocol (Phase Genomics, USA) and sequenced on the Illumina NovaSeq 6000 platform to generate 150-bp paired-end reads. HiFi reads from each of the 38 *Capsicum* accessions were assembled into contigs using Hifiasm (Cheng et al., 2021) (v0.19.6-r595) with default parameters. Hi-C reads were processed to remove adapters and low-quality sequences using Trimmomatic (Bolger et al., 2014) (v0.39) and aligned to the assembled contigs using BWA-MEM (Li, 2013) (v0.7.17) with default parameters. Chromosome-scale assemblies were generated by scaffolding the contigs with Hi-C data using Juicer (Durand et al., 2016) (v1.6) and 3D-DNA (Dudchenko et al., 2017) (v180419), followed by manual curation. Chromosome anchoring and scaffolding were validated using HiCExplorer (Wolff et al., 2022) (v3.0) and whole-genome alignments generated with MUMmer4 (Marçais et al., 2018) (v4.0.0rc1). Genome completeness was assessed using BUSCO (Simão et al., 2015) (v5.3.2) with the embryophyta_odb10 database. Base-level assembly accuracy was evaluated using Merqury (Rhie et al., 2020) (v1.3) with HiFi reads. Assembly quality was further evaluated using the LTR Assembly Index (LAI) (Ou et al., 2018) (v2.9.7).

### Repetitive sequence annotation and protein-coding gene prediction

Repetitive sequences in the genome assemblies were annotated using EDTA (Ou et al., 2019) (v2.0.1) with parameters ‘--sensitive 1 --anno 1’ and RepeatMasker (Tarailo-Graovac and Chen, 2009) (v4.1.5) with default parameters. Protein-coding genes were predicted from the repeat-masked genomes by integrating *ab initio*, homology-based and transcriptome-based evidence using the MAKER pipeline (Cantarel et al., 2008) (v3.01.04). *Ab initio* gene predictions were performed using AUGUSTUS (Stanke et al., 2006) (v3.4.0), SNAP (Korf, 2004) (v2006-07-28) and GeneMark-ES (Lomsadze et al., 2005) (v4.69_lic). Homology-based evidence included protein sequences from published pepper genomes. For transcriptome-based evidence, PacBio Iso-Seq long-read sequencing was performed on pooled fruit, leaf, flower and stem tissues. Iso-Seq reads were processed to remove primer sequences using lima (v2.12.0; https://lima.how/), clustered and polished using the IsoSeq3 pipeline (https://github.com/ylipacbio/IsoSeq3), and aligned to the corresponding genomes using minimap2 (Li, 2018) (v2.30-r1287).

To further improve gene prediction across both newly assembled and previously published *Capsicum* genomes, predicted gene models from all other genomes of the same *Capsicum* species were projected onto each target genome using Liftoff (Shumate and Salzberg, 2021) (v1.6.3) with parameters ‘-copies -sc 0.97 -exclude_partial -s 0.97 -a 0.95 -p 50’. Original and projected gene models were then integrated using EVidenceModeler (Haas et al., 2008) (v1.1.1) to generate the final gene set for each genome.

Functional annotation of the predicted protein-coding genes was performed by aligning protein sequences against the NCBI non-redundant (nr), UniProtKB/TrEMBL, UniProtKB/SwissProt (Boeckmann et al., 2003) and InterPro (Mitchell et al., 2015) databases. Gene Ontology (GO) terms were assigned using Blast2GO (Conesa and Götz, 2008) (v3.2.4).

### Gene-based pangenome construction

Gene families across the 48 *Capsicum* genomes were identified using OrthoFinder (Emms and Kelly, 2019) (v2.5.4). Pairwise syntenic gene blocks were identified among the 48 genomes using MCScanX (Wang et al., 2012) (v1.0.0), and synteny information was used to distinguish paralogous genes that were not located within the same syntenic regions. Synonymous substitution rates (*Ks*) between syntenic gene pairs were calculated using MCScanX. The median *K*_s_ value (*K*_s__median) for each syntenic block was used to exclude ancient duplications that predated *Capsicum* speciation. A maximum *K*_s__median cutoff of 0.344 was determined as the mean plus three standard deviations of all *K*_s__median values for syntenic blocks between *C. rhomboideum* and other *Capsicum* species. Pangenes were classified into four categories based on their distribution across the 48 genomes: core (present in all 48 genomes), soft-core (present in 43–47 genomes), shell (present in 2–42 genomes), and private (present in a single genome).

To reconstruct the species phylogeny, protein sequences from representative genomes of different *Capsicum* species, together with those from eggplant (*Solanum melongena* HQ-1315) (Wei et al., 2020), tomato (*S. lycopersicum* Heinz v4.0) (Hosmani et al., 2019), and potato (*S. tuberosum* DM v8.1) (Yang et al., 2023), were clustered into orthogroups using OrthoFinder (Emms and Kelly, 2019) (v2.5.4). A total of 5,020 single-copy orthologous genes were identified, and the protein sequences of each orthogroup were aligned using MUSCLE (Edgar, 2022)(v5.1). The resulting alignments were concatenated, and a maximum-likelihood species tree was reconstructed using IQ-TREE (Minh et al., 2020)(v2.4.0). Divergence times were estimated using the approximate-likelihood method implemented in MCMCtree in the PAML package (Yang, 2007) (v4.10.10), with the species-tree topology fixed. Two secondary calibration constraints based on previously reported divergence-time estimates were applied: 6.9–9.0 Ma for the divergence between *S. tuberosum* and *S. lycopersicum* and 16.1–22.7 Ma for the divergence between *Solanum* and *Capsicum* (https://timetree.org/) (Kumar et al., 2022).

### Pan-NLRome construction

NLR genes were identified from the 48 *Capsicum* genome assemblies using NLR-Annotator (Steuernagel et al., 2020) (v2.0). To validate the predicted NLR candidates, the hmmscan program from HMMER (Johnson et al., 2010) (v3.0) was used to identify the conserved NB-ARC domain (Pfam: PF00931) in the predicted protein sequences, using an E-value cutoff of 1e−3. Protein domains were further annotated using InterProScan (Jones et al., 2014) (v5.71-102.0) with the parameters ‘-dp -iprlookup -appl Pfam,Coils’. To improve coiled-coil (CC) domain prediction, DeepCoil (Ludwiczak et al., 2019) was used for secondary structure validation. Following the classification framework established in *Arabidopsis* (Van de Weyer et al., 2019), NLR proteins were categorized into four subclasses according to their domain composition: TNLs (containing at least one TIR domain), CNLs (containing at least one CC domain), RNLs (containing at least one RPW8 domain), and NLs (containing NB and LRR domain only). The predicted NLRs from the 48 genomes were clustered into orthogroups using OrthoFinder (Emms and Kelly, 2019) (v2.5.4) to construct the pan-NLRome.

### Graph-based pangenome construction and variant genotyping

A graph-based super-pangenome was constructed from 46 reference-quality *Capsicum* genome assemblies using the Minigraph-Cactus pipeline (Hickey et al., 2024), excluding two highly divergent *C. rhomboideum* genomes to reduce alignment ambiguity. The genome of *C. annuum* var. *annuum* cultivar ‘Zhudachang’ was used as the initial backbone, and the remaining assemblies were incorporated sequentially into the graph. SVs (≥ 20 bp) were extracted from the graph.

SVs identified from the graph pangenome were genotyped in the 500 core accessions (Liu et al., 2023) using PanGenie (Ebler et al., 2022) (v3.0.2) with cleaned short-read resequencing data. To assess genotyping accuracy, 35 accessions for which both chromosome-scale assemblies and short-read resequencing data were available were used as a validation dataset. Graph-based SV genotypes inferred from short-read data were compared with assembly-derived SV calls, and precision, recall, and F1 scores were calculated for each variant. SVs with F1 scores greater than 0.7 were retained for downstream population analyses. Allele frequencies inferred from graph-based short-read genotyping were further compared with assembly-derived allele frequencies to evaluate genotyping consistency.

Previously identified high-quality SNPs from the 500 core accessions, originally called against the ‘Zhangshugang’ genome (Liu et al., 2023), were lifted over to the ‘Zhudachang’ genome using whole-genome alignments generated with minimap2 (Li, 2018) (v2.30-r1287). SNP coordinates were converted based on the genome-to-genome alignments, resulting in 98,175,412 high-confidence SNPs mapped to the ‘Zhudachang’ reference genome.

### Population genomic analyses

To identify SVs differentiated between wild and domesticated populations, two independent domestication trajectories were analyzed separately: *C. baccatum* var. *baccatum* versus *C. baccatum* var. *pendulum*, and *C. annuum* var. *glabriusculum* versus *C. annuum* var. *annuum*. For each SV, allele frequencies were calculated within each population, and differences between wild and domesticated groups were evaluated using Fisher’s exact test. The resulting *P* values were adjusted for multiple testing using the Benjamini–Hochberg false discovery rate (FDR) procedure (Benjamini and Hochberg, 1995). SVs with an FDR < 0.001 and a greater than twofold difference in allele frequency were considered differentiated between wild and domesticated populations.

To infer the unfolded site frequency spectrum (SFS) and quantify genetic burden, ancestral and derived states of SNPs and SVs were inferred using *C. pubescens* as the reference for ancestral-state polarization. For each variant, the major allele in the *C. pubescens* population was designated as ancestral, whereas the alternative allele was considered derived. Unfolded SFSs were calculated separately for SNPs and SVs based on derived allele frequencies. For each accession, SNP and SV burdens were calculated as the number of derived nonsynonymous SNPs or SVs, respectively, normalized by the number of derived synonymous SNP alleles.

### Plant growth and phenotyping

A field trial involving 500 core *Capsicum* accessions was conducted in Damao Village, Jiyang District, Sanya, China (18.23°N, 109.52°E). Seeds were sown on September 20, 2022, and seedlings were transplanted on October 22, 2022. The trial concluded on April 10, 2023. Plants were grown in sandy loam soil under a 40-mesh nylon net. Morphological traits were recorded for accessions that successfully produced fruits. A second field trial involving the same 500 accessions was conducted at the same location during the 2023–2024 growing season for fruit metabolite analyses. Only accessions yielding sufficient fruit material (>500 g) were included in the capsaicinoid, carotenoid, and volatile compound analyses. Fruits were washed, homogenized, freeze-dried for 72 h, ground into powder, and passed through a 40-mesh sieve before metabolite analysis.

### Capsaicinoid quantification

For capsaicinoid analysis, 0.25 g of fruit powder was mixed with 15 mL of methanol/water (7:3, v/v) containing 0.03% butylated hydroxytoluene (BHT; Sigma-Aldrich). The mixture was vortexed for 1 min, ultrasonicated for 30 min, and centrifuged at 10,000 rpm for 5 min. The supernatant was collected, diluted 200-fold, and filtered through a 0.22-μm membrane filter. Capsaicinoids were quantified using a Thermo Scientific Ultimate 3000 UHPLC system coupled to a TSQ Endura triple-quadrupole mass spectrometer equipped with an electrospray ionization (ESI) source. Chromatographic separation was performed on a Waters XBridge Peptide BEH C18 column (100 mm × 2.1 mm, 3.5 μm) maintained at 35 °C, using 0.05% acetic acid in water (A) and acetonitrile (B) as the mobile phase. The gradient increased from 50% to 80% B over 7 min, followed by column re-equilibration. The flow rate was 0.4 mL/min, and the injection volume was 5 μL. Detection was carried out in multiple reaction monitoring (MRM) mode with the following transitions: nordihydrocapsaicin ([M+H]+, 294.1→137.1, CE 10 V), capsaicin ([M +H]+, 306.1→137.1, CE 14 V), dihydrocapsaicin ([M+H]+, 308.2→137.1, CE 11 V), and homodihydrocapsaicin ([M-H]-, 320.2→184.1, CE 15 V).

### Carotenoid quantification

For carotenoid analysis, 0.2 g of pepper fruit powder was mixed with 30 mL of methanol-diethyl ether (6:4, v/v) containing 0.03% BHT. The mixture was vortexed for 1 min, sonicated at 25 °C for 30 min, and centrifuged at 10,000 rpm for 5 min. The supernatant was filtered through a 0.22-μm membrane filter. Carotenoids were quantified using the same UHPLC system described above. Chromatographic separation was performed on a YMC Carotenoid column (100 mm × 2.0 mm, 3 μm) maintained at 35 °C. The mobile phases consisted of 0.01% formic acid with 5 mmol/L ammonium formate in water (A) and methyl tert-butyl ether containing 0.01% BHT (B). The gradient was programmed as follows: 0-3 min, 10-15% B; 3-5 min, 15-80% B; 5-8 min, 80-100% B; 8-10 min, 100% B; 10-10.1 min, 100-10% B; and 10.1-15 min, 10% B. The flow rate was 0.3 mL/min, and the injection volume was 5 μL. Detection was performed in positive ionization mode with MRM. Optimized parameters were as follows: ESI source temperature, 325 °C; ion transfer tube temperature, 325 °C; spray voltage, 3.6 kV; sheath gas, 30 arbitrary units (Arb); auxiliary gas, 10 Arb; and sweep gas, 0 Arb.

### Volatile profiling

For volatile analysis, approximately 3 g of ripe pepper fruit powder was placed in a 20-mL headspace vial, and 10 µL of 2-octanol (200 µg/mL in distilled water) was added as an internal standard. Samples were equilibrated at 40 °C for 30 min, and volatile compounds were extracted at 35 °C for 20 min using a solid-phase microextraction (SPME) fiber (DVB/C-WR/PDMS, 10 mm, 50/30 µm; GERSTEL GmbH & Co. KG, Mülheim, Germany). The fiber was thermally desorbed in the gas chromatograph (GC) injection port for 5 min under splitless conditions. Volatile compounds were analyzed using an Agilent 7890A GC coupled to an Agilent 7200 QTOF mass spectrometer (Agilent Technologies Inc., Santa Clara, CA, USA) equipped with a DB-5MS capillary column (30 mm × 0.32 mm, 0.25 µm; Agilent Technologies). The temperature program was as follows: 40 °C for 1 min, increased at 2 °C/min to 120 °C (held for 3 min), then at 10 °C/min to 160 °C (held for 2 min), and finally at 20 °C/min to 240 °C. Helium (≥99.999%) was used as the carrier gas at a flow rate of 1.0 mL/min. Mass spectra were acquired in electron ionization (EI) mode at 70 eV, with the transfer line and ion source temperatures set to 250 °C and 230 °C, respectively.

### Genome-wide association studies

Genome-wide association studies (GWAS) were performed separately for SNPs and SVs using EMMAX (Kang et al., 2010). The top five principal components and the Balding–Nichols (BN) kinship matrix were included as covariates to account for population structure and genetic relatedness. Genome-wide significance thresholds were determined based on the effective numbers of independent SNPs and SVs, estimated separately using the Genetic Type I error Calculator (Li et al., 2012). Candidate association regions were defined based on linkage disequilibrium (LD; *r*^2^ ≥ 0.8) between the lead variant and neighboring variants.

### RNA-seq experiments and data analysis

*C. annuum* var. *annuum* cultivar ‘Zhangshugang’ and two non-pungent mutant lines (c68 and ss97) were used for RNA-seq analysis. Fruit placenta and pericarp tissues were collected at 15, 20, and either 25 or 30 days after pollination (DAP), with three biological replicates per sample. Additional raw RNA-seq datasets were obtained from previous studies (Liu et al., 2023; Chen et al., 2024). Adapters and low-quality sequences were removed using Trimmomatic (Bolger et al., 2014) (v0.39) with the parameters ‘TruSeq3-PE-2.fa:2:30:10:1:TRUE SLIDINGWINDOW:4:20 LEADING:3 TRAILING:3 MINLEN:40’. Poly(A) tails were trimmed using prinseq++ (Cantu et al., 2019) (v1.2), and rRNA-derived reads were removed by mapping against an rRNA reference database (Chuvochina et al., 2026). The cleaned reads were then aligned to the ‘Zhudachang’ genome using HISAT2 (Kim et al., 2019) (v 2.2.1) with default parameters. Gene-level read counts were generated using featureCounts (Liao et al., 2014) (v2.1.1) and normalized to transcripts per million (TPM). Genes differentially expressed between ‘Zhangshugang’ and each of the two non-pungent mutant lines were identified separately using DESeq2 (Love et al., 2014) (v1.52.0) based on raw read counts, with thresholds of |log2(fold change)| ≥ 1 and an adjusted *P* value < 0.05.

### Virus-induced gene silencing

VIGS was performed as previously described (Zhou et al., 2021; Liu et al., 2026), with minor modifications. A 300-bp *RAP2*-specific fragment, predicted to have no off-target effects using the VIGS Tool (Fernandez-Pozo et al., 2015), was cloned into the pTRV2 vector. The resulting construct was introduced into competent cells of *Agrobacterium tumefaciens* strain GV3101 using the freeze–thaw method, and positive clones were verified by colony PCR. Positive bacterial cultures were incubated overnight in Luria–Bertani (LB) medium supplemented with 50 mg/L rifampicin and 50 mg/L kanamycin at 28 °C with shaking at 200 rpm. Cells were harvested by centrifugation at room temperature and resuspended in infiltration buffer (10 mM MgCl₂, 10 mM MES, pH 5.6, and 200 μM acetosyringone) to a final OD₆₀₀ of 1.0. Equal volumes of *A. tumefaciens* cultures harboring pTRV1 and either pTRV2-*RAP2*, empty pTRV2, or pTRV2-*PDS* were mixed and incubated at 28 °C in the dark for 4 h. Pepper seedlings with fully expanded cotyledons but no true leaves (approximately 14 days after sowing) were subjected to water deprivation for 2 days. The *Agrobacterium* mixtures were then infiltrated into the abaxial side of the cotyledons using a needleless syringe (approximately 10 μL per cotyledon) until the cotyledons became water-soaked. Following infiltration, plants were maintained in darkness at 22 °C for 24 h and then transferred to a controlled-environment growth chamber (22 °C day/18 °C night, 65% relative humidity, 16 h light/8 h dark, and a light intensity of 150 μmol m⁻² s⁻¹). Mixtures containing pTRV1 and either empty pTRV2 or pTRV2-*PDS* were used as negative and positive controls, respectively. Each treatment included 20 plants, and the experiment was independently repeated three times. Four weeks after infiltration, newly developed leaves were collected for RT-qPCR analysis to access *RAP2* silencing efficiency. Plants with confirmed silencing were maintained until fruit ripening, and ripe fruits were collected for RT-qPCR analysis, phenotypic characterization, and carotenoid profiling.

### Genomic prediction

Genomic prediction was conducted separately for each trait using either SNP-only or combined SNP+SV variant sets. Variant filtering was performed independently for each trait using accessions with available phenotypic data. Within each trait-specific subset, variants with a genotype missing rate >20%, minor allele count <10, or minor allele frequency <0.05 were excluded. Variants were further pruned based on pairwise linkage disequilibrium (LD) using an *r^2^* threshold of 0.1. The resulting genotype matrices were used for trait-specific marker selection and genomic prediction using CropGBM (Yan et al., 2021) (v1.1.2). Prediction accuracy was evaluated using 50 repetitions of five-fold cross-validation and measured as the Pearson correlation coefficient between observed and predicted phenotypic values.

## Data availability

Raw HiFi, Hi-C and RNA-seq reads have been deposited in the NCBI BioProject database under accession number PRJNA1026847. Genome assemblies have been deposited in the NCBI BioProject database under accession number PRJNA1007743. Genome assemblies and annotations, and variant files in VCF format are available through the Pepper Genomics Database (http://ted.bti.cornell.edu/ftp/pepper/graph_pan).

## Acknowledgements

The authors thank Jing Han and Bosheng Li from Peking University Institute of Advanced Agricultural Sciences for providing spatial expression data of *CS-1*, *CS-2,* and *pAMT5*. This research was supported by Major International (Regional) Joint Research Project (32320103011), Key Program of the National Natural Science Foundation of China (32130097), Yuelushan Laboratory Breeding Program (YLS-2025-ZY01001), and Major Program of the National Natural Science Foundation of China (32494784).

## Author contributions

F.L., Z.F., and X.Zou designed and managed the project. F.L. and R.J. coordinated genome sequencing. J.Z., X.Zhao, and H.S. performed genomic data analyses. F.Z., X.D., L.O., and L.Y. conducted field trials and plant cultivation. Y.Li, H.X., S.Y., Z.L., and B.Y. performed phenotypic evaluation. R.W., C.X., and B.H. conducted metabolite profiling and data analysis. Y.Liu collected samples for transcriptome sequencing. H.Y. and Z.W. contributed to data curation. P.Z. performed functional validation experiments. J.Z., X.Zhao, and H.S. wrote the manuscript. Z.F. revised the manuscript.

