## Supplementary Figures for "A super-pangenome of *Capsicum* reveals genomic basis of domestication, pungency, and metabolic diversity"

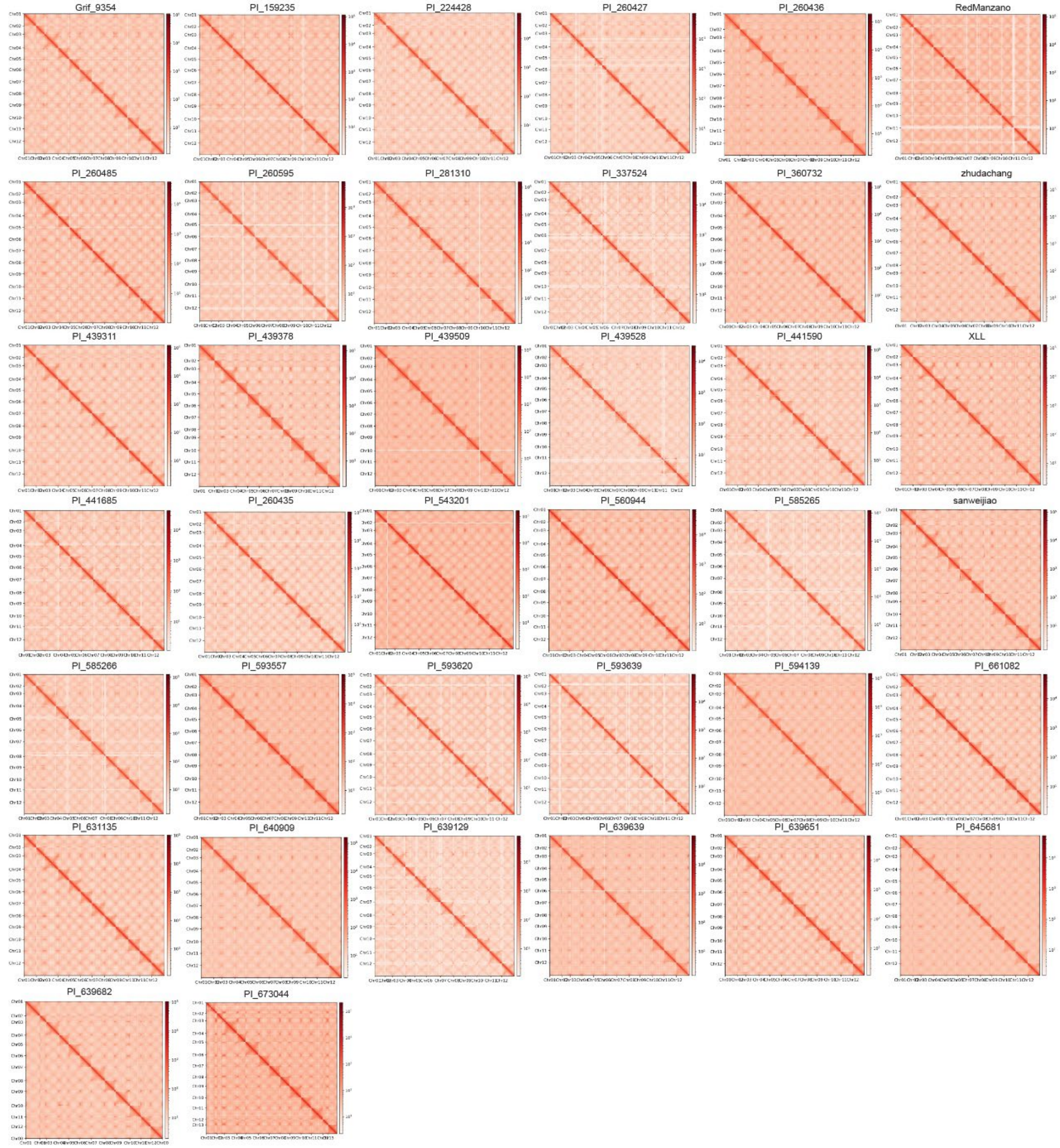


**Supplementary Fig. 1** Hi-C interaction heatmaps of the 38 newly assembled *Capsicum* genomes.

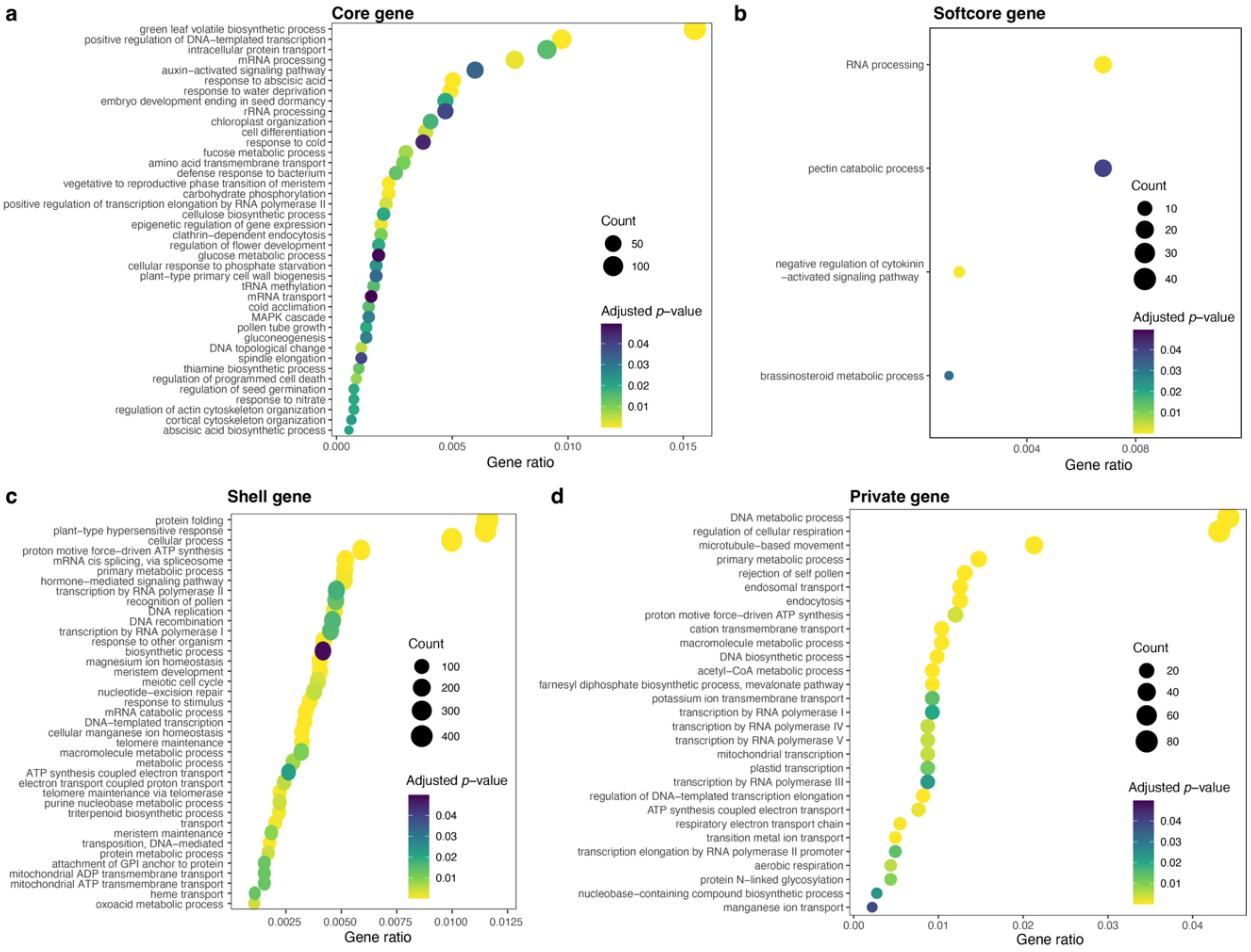


**Supplementary Fig. 2** Gene ontology (GO) term enrichment analysis of pangene categories in the *Capsicum* super-pangenome. **a-d,** Significantly enriched GO terms for core (**a**), soft-core (**b**), shell (**c**), and private (**d**) pangenes.

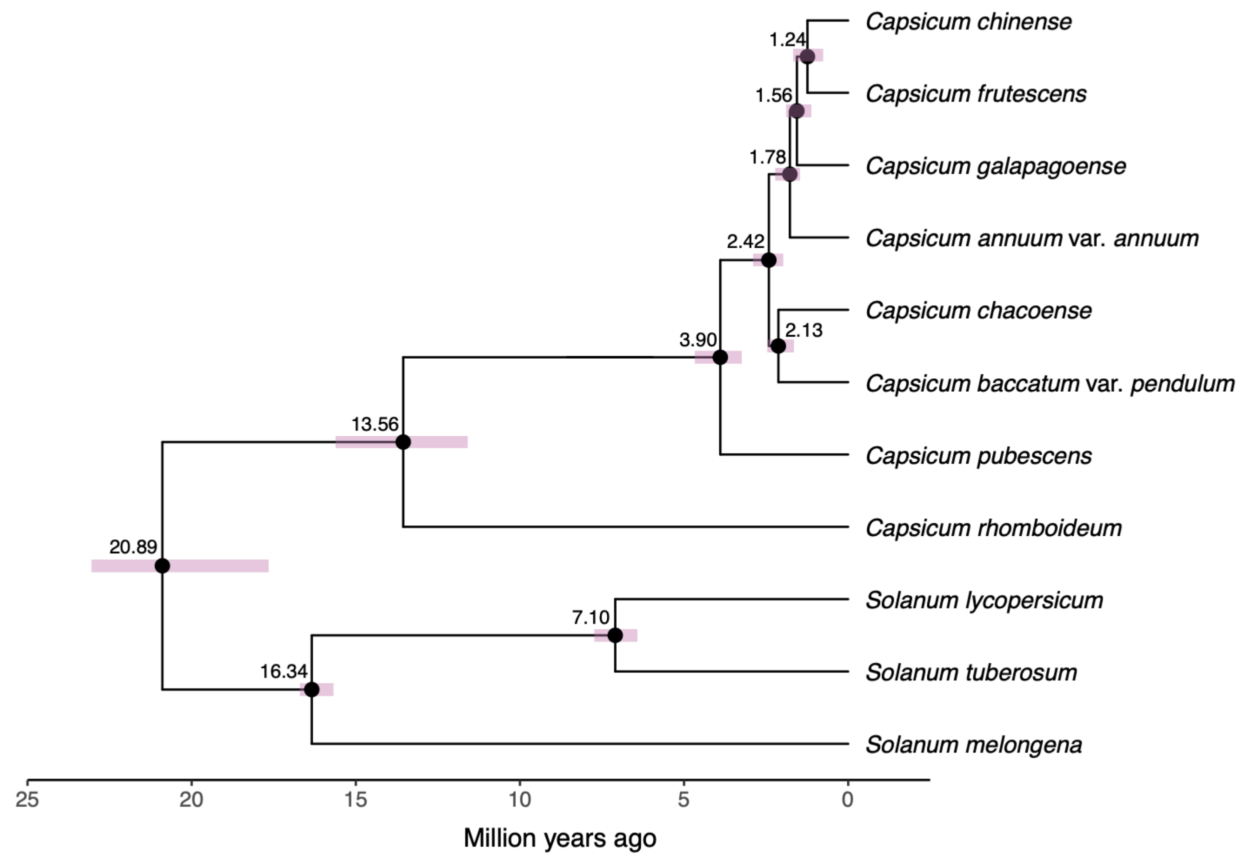


**Supplementary Fig. 3** Divergence times among major *Capsicum* lineages and representative *Solanum* crops (tomato, potato and eggplant). The time-calibrated phylogeny was inferred using single-copy orthologs. Numbers at internal nodes indicate estimated divergence times in millions of years ago (Ma), and horizontal shaded bars represent the 95% highest posterior density (HPD) intervals of the divergence-time estimates.

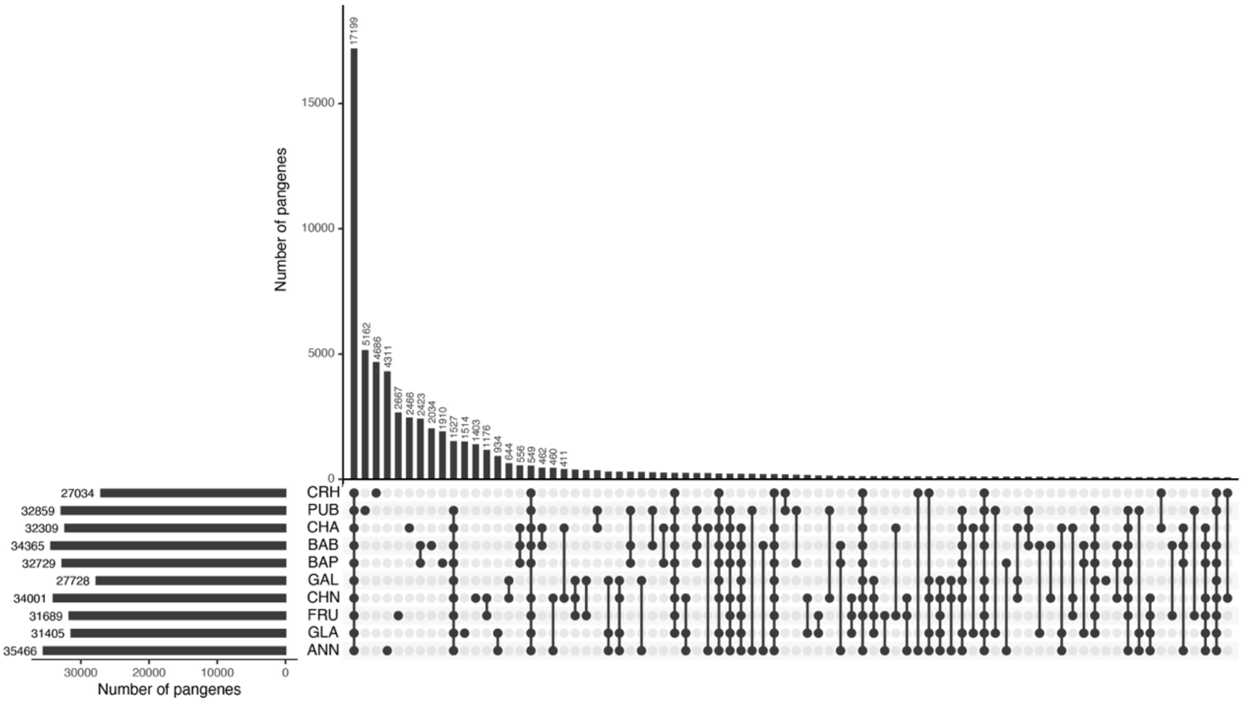


**Supplementary Fig. 4** Presence-absence variation (PAV) of pangenes across major *Capsicum* groups. The UpSet plot shows the top 75 pangene intersections among ten *Capsicum* groups. Horizontal bars indicate the total number of pangenes in each group, and vertical bars indicate the number of pangenes in each intersection, with the sizes of the top 20 intersections shown above the corresponding bars. CRH, *C. rhomboideum*; PUB, *C. pubescens*; CHA, *C. chacoense*; BAB, *C. baccatum* var. *baccatum*; BAP, *C. baccatum* var. *pendulum*; GAL, *C. galapagoense*; CHN, *C. chinense*; FRU, *C. frutescens*; GLA, *C. annuum* var. *glabriusculum*; ANN, *C. annuum* var. *annuum*.

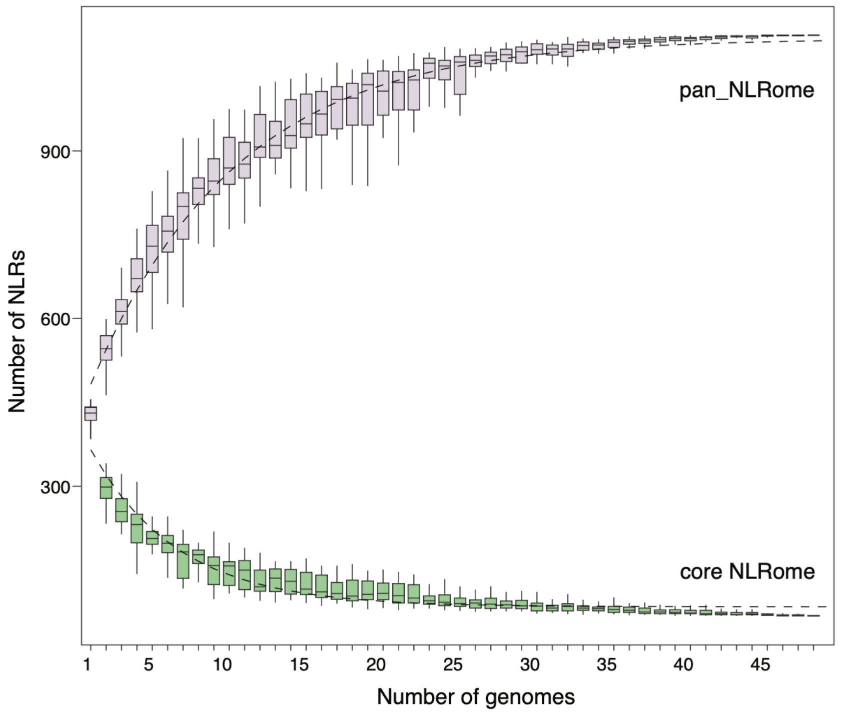


**Supplementary Fig. 5** Pan-NLRome expansion and core NLRome contraction with increasing number of *Capsicum* genomes. For each given number of accessions, 100 random combinations of accessions were sampled. Boxplots show the distributions across random combinations, and dashed lines indicate the fitted accumulation and contraction curves for the pan-NLRome and core NLRome, respectively. For each boxplot, the lower and upper bounds indicate the first and third quartiles, respectively, the centerline indicates the median and the whiskers extend to 1.5× interquartile range.

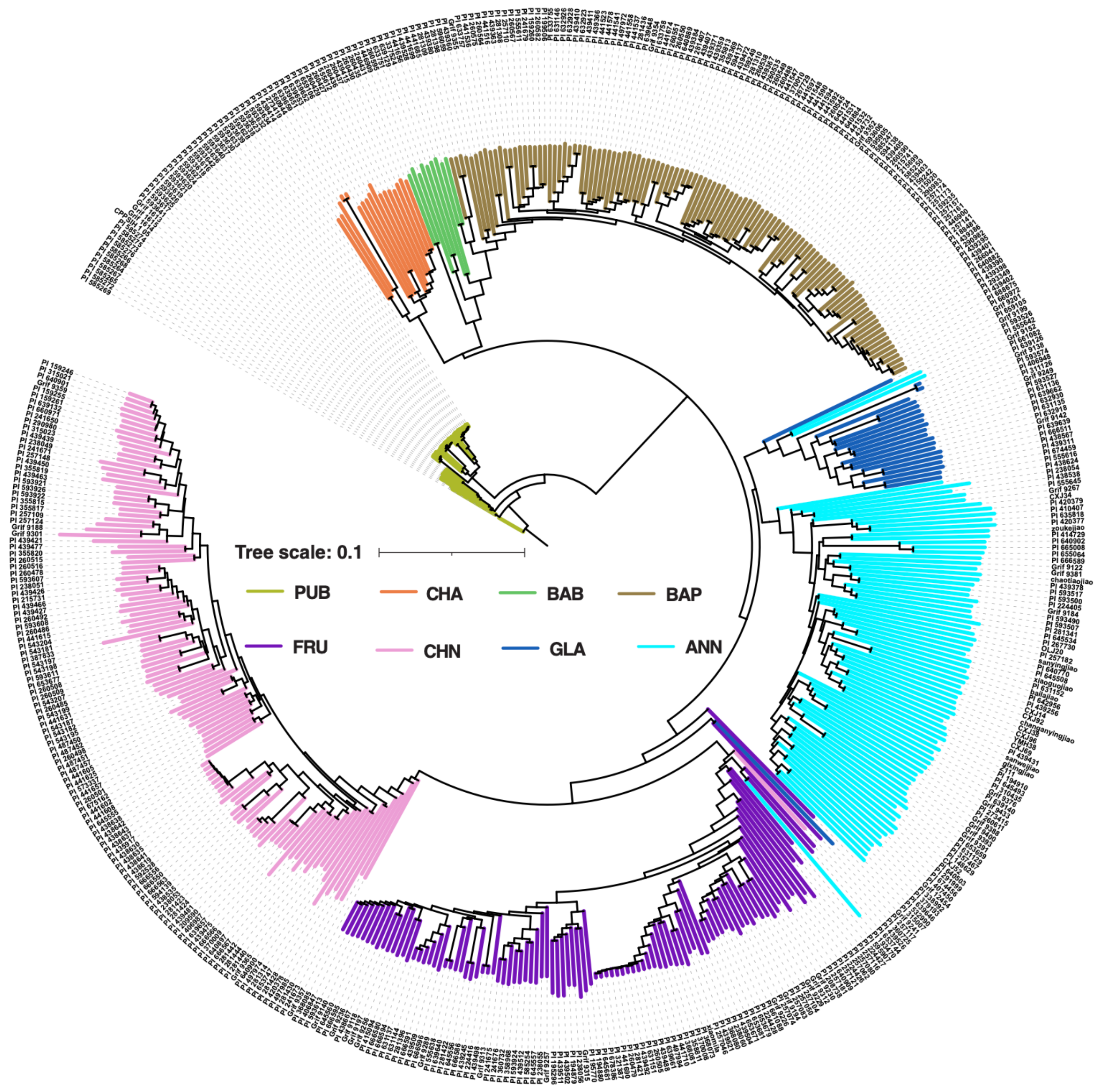


**Supplementary Fig. 6** Phylogenetic relationships of *Capsicum* accessions based on SVs. Branch colors indicate the *Capsicum* groups to which the accessions belong. PUB, *C. pubescens*; CHA, *C. chacoense*; BAB, *C. baccatum* var. *baccatum*; BAP, *C. baccatum* var. *pendulum*; FRU, *C. frutescens*; CHN, *C. chinense*; GLA, *C. annuum* var. *glabriusculum*; ANN, *C. annuum* var. *annuum*.

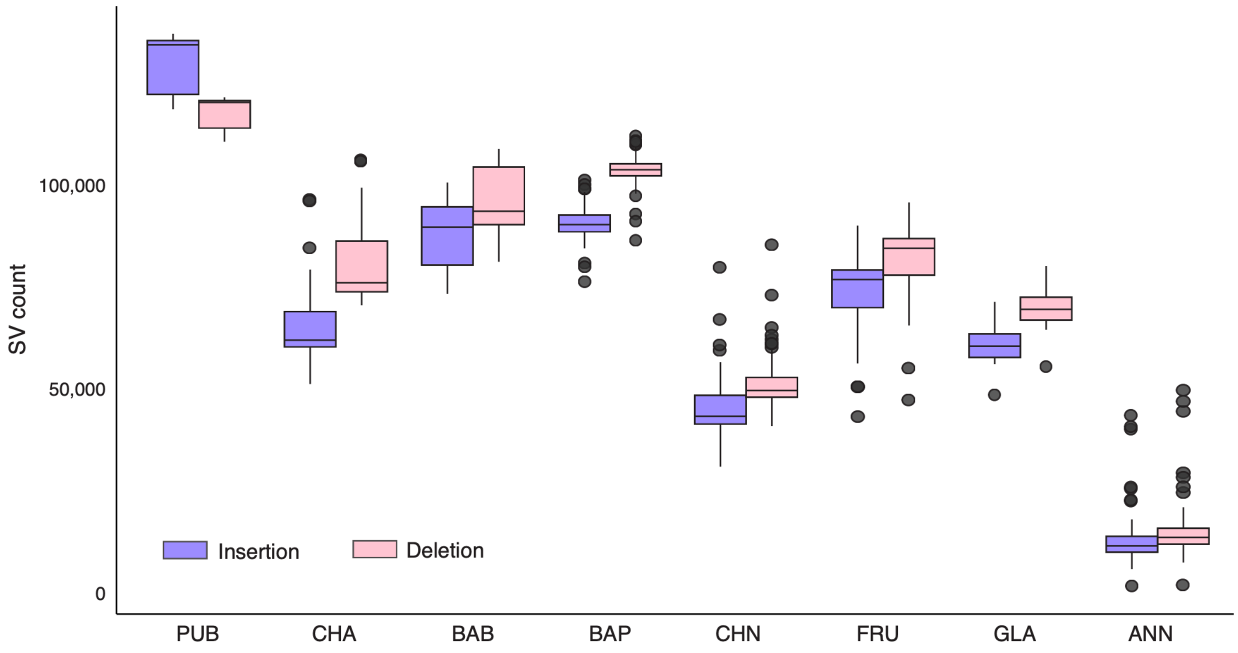


**Supplementary Fig. 7** Distribution of insertion and deletion counts per accession across eight *Capsicum* groups. For each boxplot, the lower and upper bounds indicate the first and third quartiles, respectively, the centerline indicates the median and the whiskers extend to 1.5× interquartile range. PUB, *C. pubescens*; CHA, *C. chacoense*; BAB, *C. baccatum* var. *baccatum*; BAP, *C. baccatum* var. *pendulum*; CHN, *C. chinense*; FRU, *C. frutescens*; GLA, *C. annuum* var. *glabriusculum*; ANN, *C. annuum* var. *annuum*.

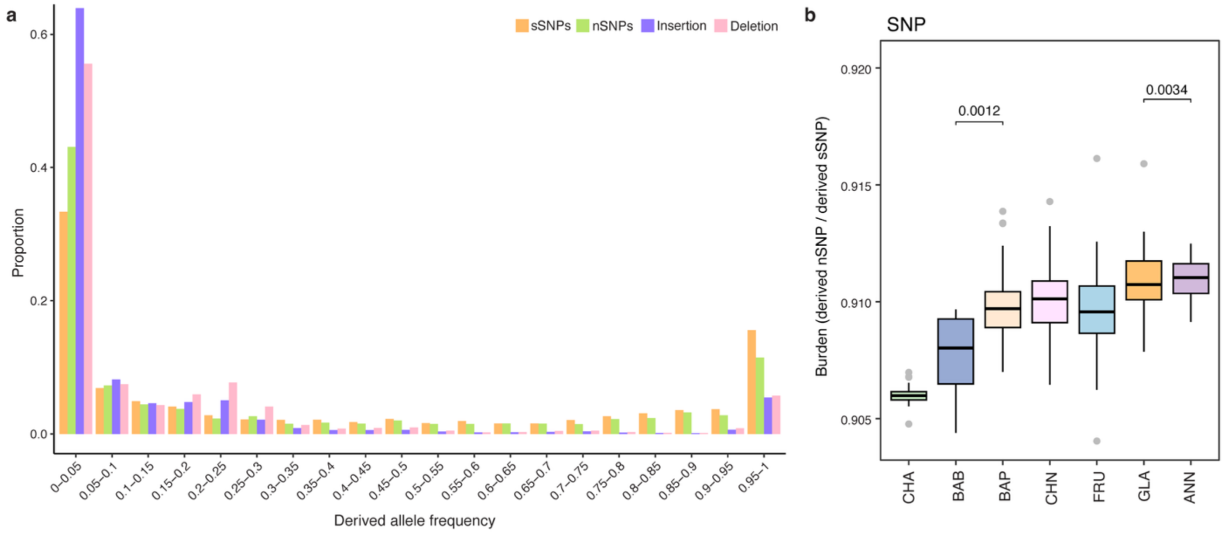


**Supplementary Fig. 8** Derived allele frequency distributions and nonsynonymous SNP burdens across Capsicum populations. (**a**) Distribution of derived allele frequencies for synonymous SNPs (sSNPs), nonsynonymous SNPs (nSNPs), insertions, and deletions in *Capsicum*. (**b**) nSNP burdens across major *Capsicum* groups. Numbers above brackets indicate P values for comparisons between wild and domesticated groups (two-sided Student's *t*-tests). For each boxplot, the lower and upper bounds indicate the first and third quartiles, respectively, the centerline indicates the median and the whiskers extend to 1.5× interquartile range. CHA, *C. chacoense*; BAB, *C. baccatum* var. *baccatum*; BAP, *C. baccatum* var. *pendulum*; FRU, *C. frutescens*; CHN, *C. chinense*; GLA, *C. annuum* var. *glabriusculum*; ANN, *C. annuum* var. *annuum*.

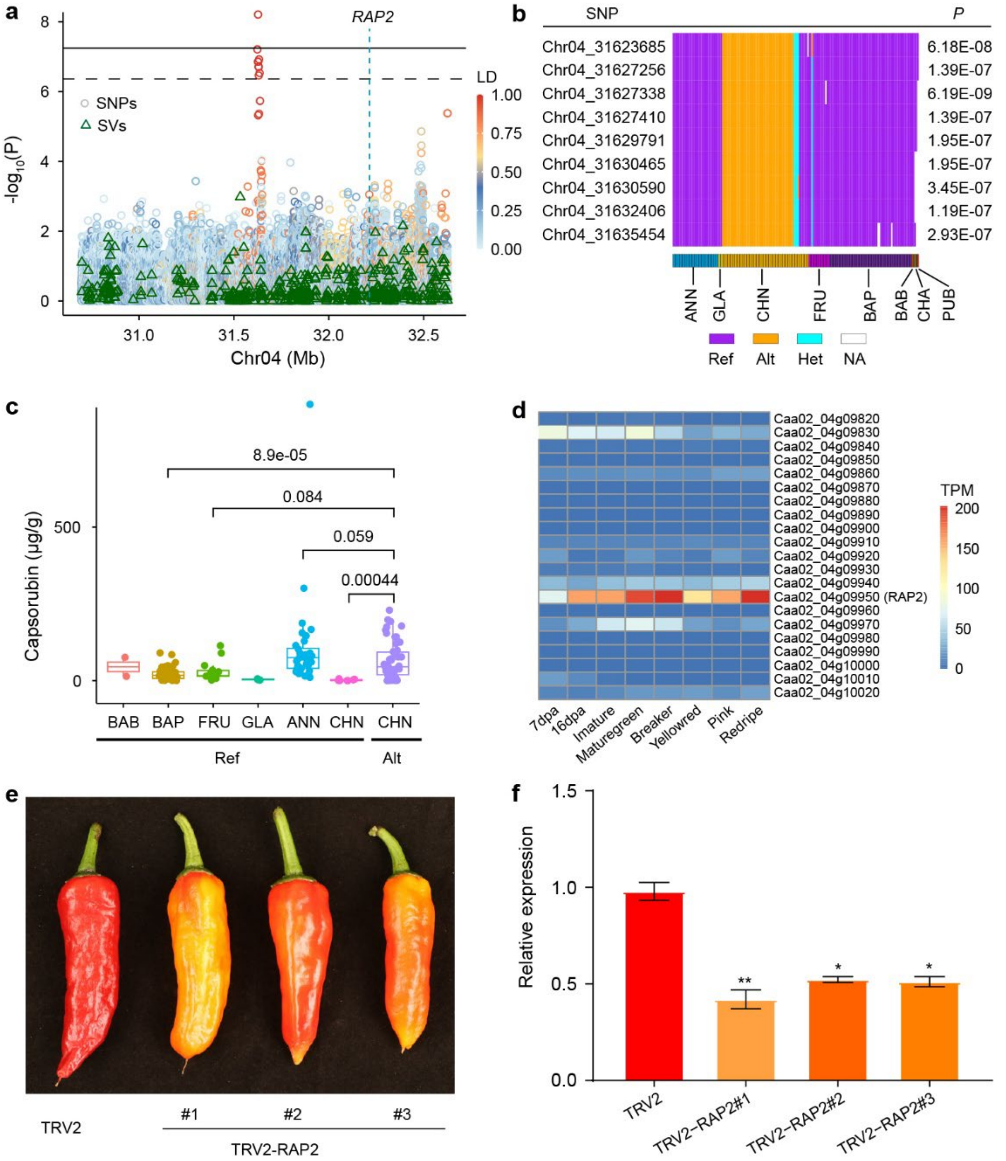


**Supplementary Fig. 9** Association mapping and functional validation identify *RAP2* as a regulator of capsorubin accumulation in pepper fruit. **a**, Regional Manhattan plot showing SNP- and SV-based associations with capsorubin content on chromosome 4. Circle colors indicate linkage disequilibrium (LD) with the lead SNP (Chr04:31,627,338). **b**, Genotype patterns of SNPs exceeding the suggestive significance threshold across different Capsicum groups, with corresponding association *P* values. Ref, reference allele; Alt, alternative allele; Het, heterozygous; NA, missing genotype. **c**, Capsorubin contents in accessions from different *Capsicum* groups carrying the reference (Ref) or alternative (Alt) allele of the lead SNP. Numbers above brackets indicate *P* values for pairwise comparisons. Boxes indicate the interquartile range, center lines indicate the median, and whiskers extend to 1.5 × the interquartile range. BAB, *C. baccatum* var. *baccatum*; BAP, *C. baccatum* var. *pendulum*; FRU, *C. frutescens*; CHN, *C. chinense*; GLA, *C.*

*annuum* var. *glabriusculum*; ANN, *C. annuum* var. *annuum*. **d**, Expression heatmap of genes within the associated genomic region across fruit developmental stages. **e**, Fruit phenotypes following virus-induced gene silencing (VIGS) of *RAP2*. TRV2, empty vector control; TRV2-RAP2, three independent *RAP2*-VIGS lines. **f**, Relative expression of *RAP2* in ripe fruits of the TRV2 control and *RAP2*-silenced lines, determined by qRT–PCR. Expression levels were normalized to the TRV2 control. Statistical significance of difference between each *RAP2*-silenced line and the TRV2 control was determined using a two-sided Student's *t*-test comparing each silenced sample with the TRV2 control (*, $P < 0.05$; **, $P < 0.01$).

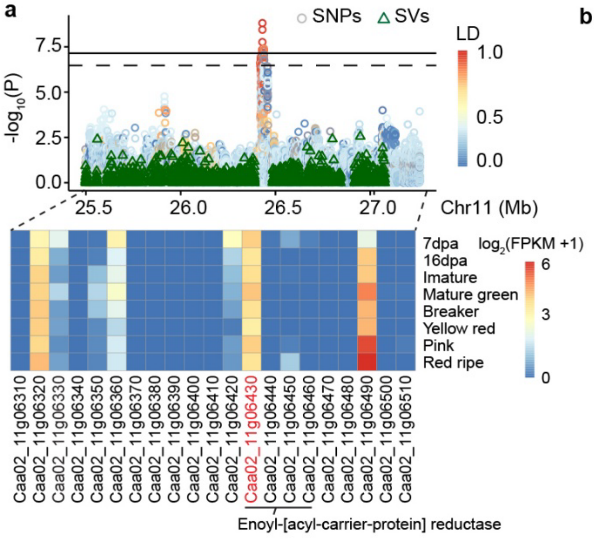


**Supplementary Fig. 10** Association mapping identifies an *ENR* gene cluster associated with capsaicinoid accumulation. **a**, Regional Manhattan plot showing SNP- and SV-based associations with homodihydrocapsaicin content on chromosome 11 and expression heatmap of genes within the associated region across fruit developmental stages. Circle colors indicate linkage disequilibrium (LD) with the lead SNP (Chr11:26,431,163). **b**, Contents of nordihydrocapsaicin, capsaicin, dihydrocapsaicin, and homodihydrocapsaicin in accessions carrying the reference (Ref), heterozygous (Het), or alternative (Alt) allele of the lead SNP. Numbers above brackets indicate *P* values for pairwise comparisons (two-sided Student's *t*-test). For each boxplot, the lower and upper bounds indicate the first and third quartiles, respectively, the centerline indicates the median and the whiskers extend to 1.5× interquartile range.

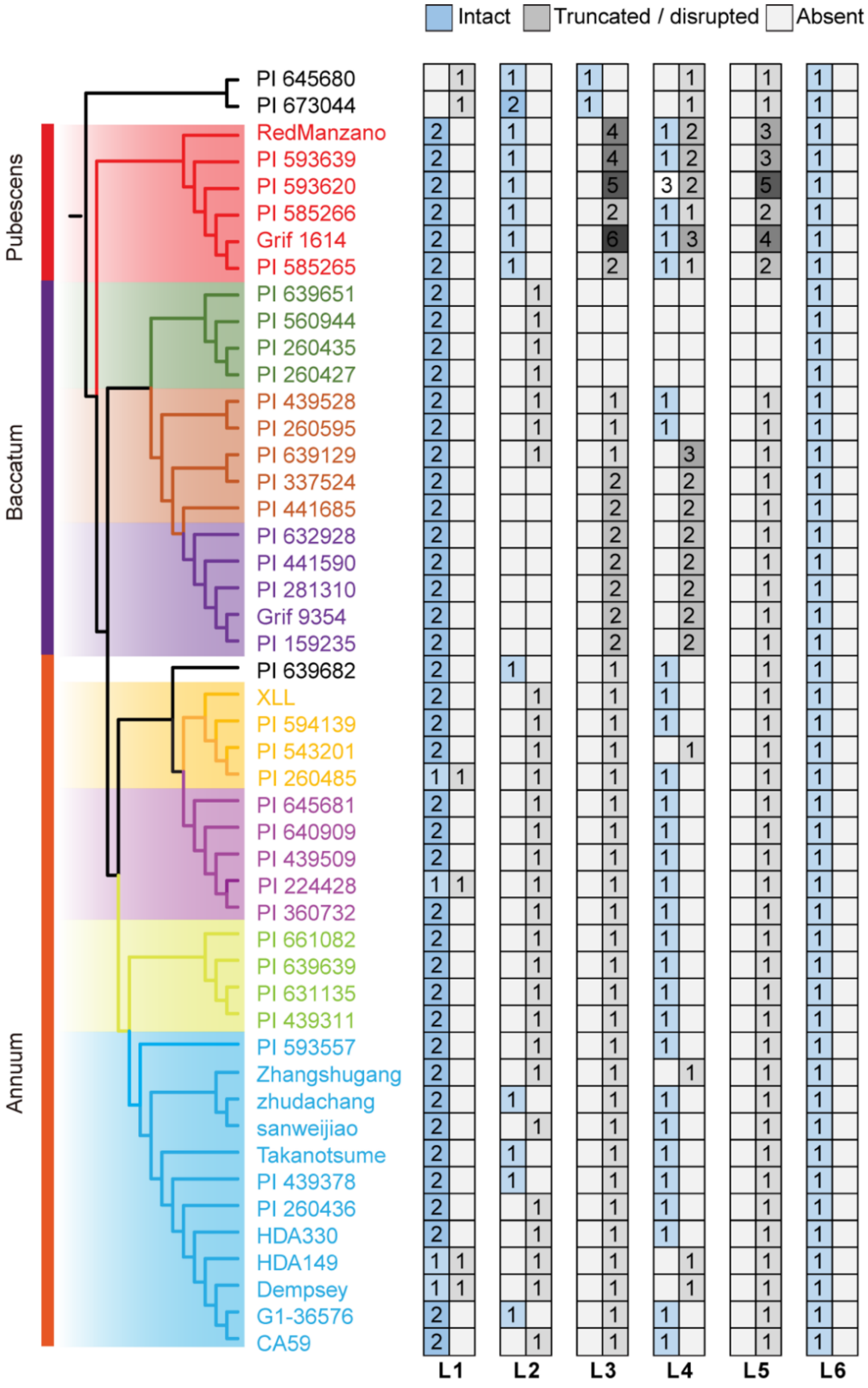


**Supplementary Fig. 11** Distribution of six *AT3* paralogous lineages at the *Pun1* locus across 48 *Capcisum* genomes. Phylogenetic relationships among the 48 *Capcisum* accessions are shown on the left. The presence and copy number of six *AT3* paralogous lineages (L1–L6) are shown on the right. Numbers within boxes indicate the number of paralogues belonging to each lineage in each genome. *CS-1* and *CS-2* belong to the lineage L1.

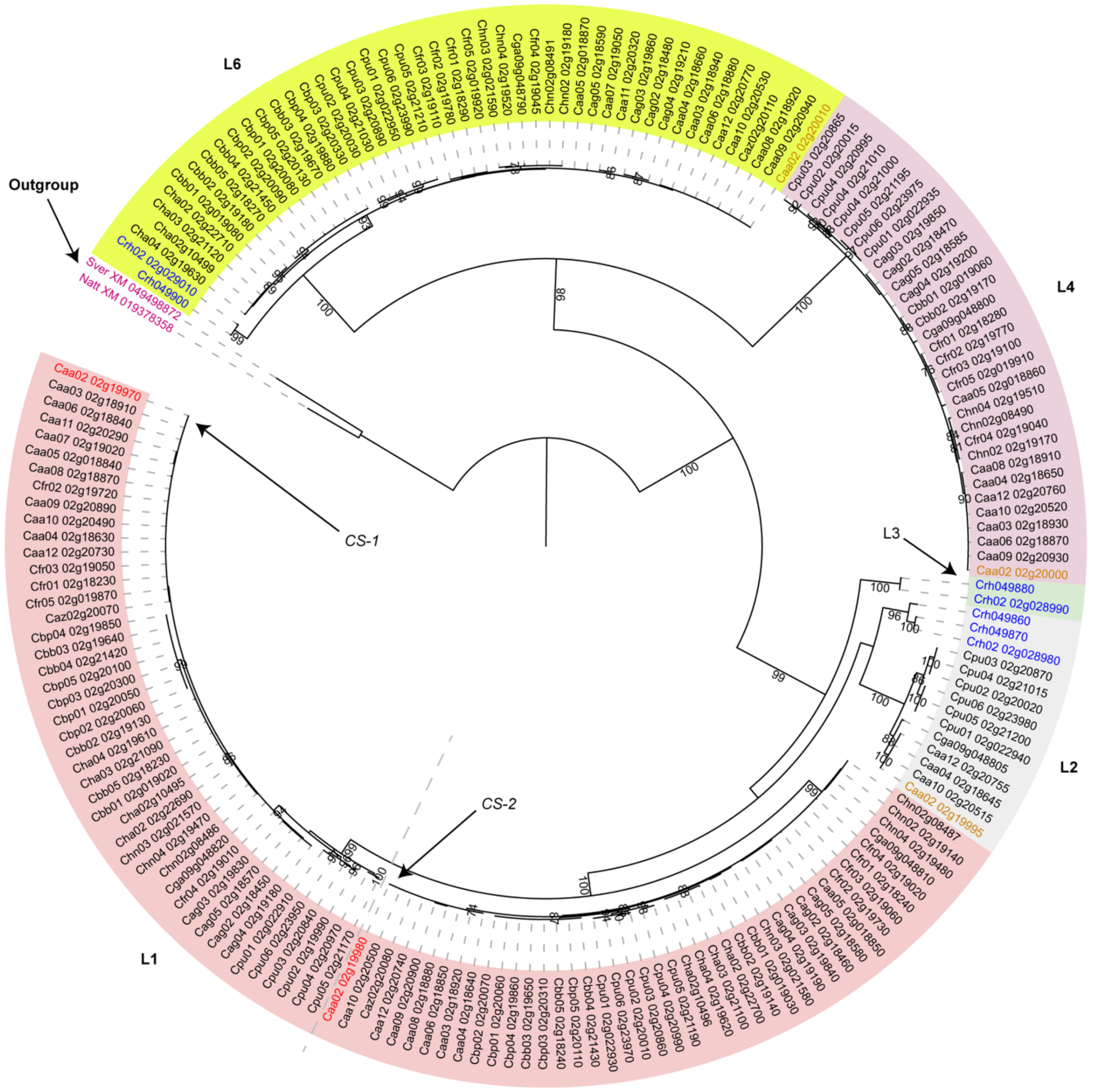


**Supplementary Fig. 12** Maximum-likelihood phylogeny of intact *AT3* paralogues. The rooted phylogenetic tree was reconstructed from coding sequences of the second exon using IQ-TREE (v3.1.1) with the substitution model selected by ModelFinder. Branch support was assessed using 1,000 standard nonparametric bootstrap replicates, with bootstrap values shown at the corresponding nodes. Five major *AT3* paralogous lineages (L1-L4 and L6) are highlighted by different background colors. Lineage L5 is not shown because no intact copy was identified among the genomes analyzed. Within L1, orthologues of *CS-1* and *CS-2* form two distinct, strongly supported clades. *CS-1* and *CS-2* from 'Zhudachang' are highlighted in red, other 'Zhudachang' paralogues in brown, *C. rhomboideum* homologues in blue, and outgroup homologues in purple.

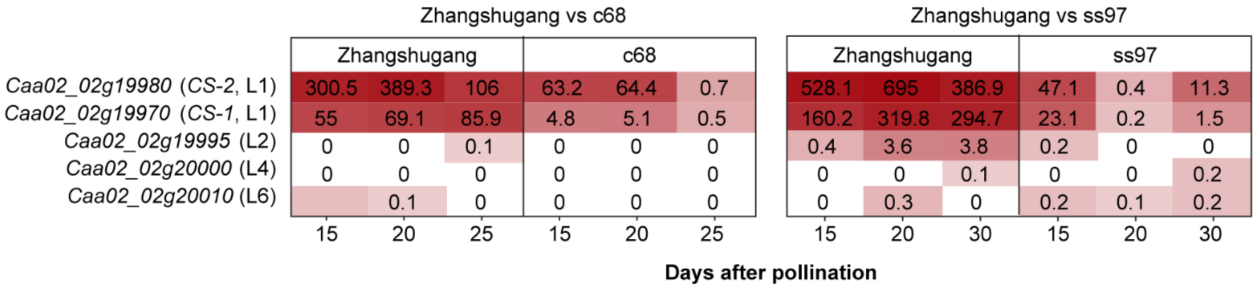


**Supplementary Fig. 13** Expression profiles of intact *AT3* paralogues at the *Pun1* locus in the placenta of the pungent cultivar 'Zhangshugang' and two non-pungent mutants, c68 and ss97. Expression profiles of five intact *AT3* paralogues identified in the 'Zhudachang' reference genome are shown. Values indicate mean transcript abundance (transcripts per million; TPM) from three biological replicates. The 'Zhangshugang'–c68 and 'Zhangshugang'–ss97 comparisons were derived from independent experimental batches.
